# The sunflower WRINKLED1 transcription factor regulates fatty acid biosynthesis genes through an AW box binding sequence with a particular base bias

**DOI:** 10.1101/2022.03.13.484140

**Authors:** R Sánchez, I González-Thuillier, M Venegas-Calerón, R Garcés, JJ Salas, E Martínez-Force

## Abstract

Sunflower (*Helianthus annuus* L.) is an important oilseed crop in which the biochemical pathways leading to seed oil synthesis and accumulation have been widely studied. However, how these pathways are regulated is less well understood. The WRINKLED1 (WRI1) transcription factor is considered a master regulator in the transcriptional control of triacylglycerol biosynthesis, acting through the AW box binding element (CNTNG(N)_7_CG) that resides in the promoter of target genes. Here, we identified the sunflower *WRI1* gene and characterized its activity in electrophoretic mobility shift assays. We studied its role as a co-regulator of sunflower genes involved in plastidial fatty acid synthesis, identifying genes bound by this transcription factor. Sunflower WRI1-targets included genes encoding all subunits of the pyruvate dehydrogenase complex, the α-CT and BCCP genes of the acetyl-CoA carboxylase complex, genes encoding acyl carrier proteins and key genes of the fatty acid synthase complex (KASIII, KASI and KAR), together with the *FATA1* gene. As such, sunflower WRI1 regulates seed plastid fatty acid biosynthesis in a coordinated manner, establishing a WRI1 push and pull strategy that drives oleic acid synthesis for its export into the cytosol. We also analyzed the sequence of the functional sunflower AW box, determining the base bias at the *N* positions in the active sunflower AW box motif. Accordingly, we conclude that the sunflower AW box is sequence-sensitive at the non-conserved positions, enabling WRI1-binding. Moreover, we found that sunflower WRI1 could bind to a non-canonical AW-box motif, opening the possibility of searching for new target genes.

## INTRODUCTION

Plants oils are important elements of human diet, as well as for industrial applications (e.g. detergents and lubricants) and biodiesel production. As the global demand for plant oils is rapidly increasing, reliance on the production of higher plant oils augments. Many plants synthesize triacylglycerol (TAG) in seeds as an essential means to store and provide energy for seedling development (Chapman and Ohlrogge, 2012). TAG biosynthesis involves two major steps, fatty acid (FA) biosynthesis and TAG assembly, in which different cell compartments participate: plastids, the cytosol and the endoplasmic reticulum (Ohlrogge and Chapman, 2011). FA and TAG synthesis is regulated by different factors in the different cellular compartments of plants, and each reaction is catalyzed by specialist enzymes (Li-Beisson et al., 2013).

The sugars derived from photosynthesis in source tissues are imported into the developing seeds and converted into FA precursors in the cytosol by glycolysis (Durrett et al., 2008; Baud and Lepiniec, 2010). *De novo* FA biosynthesis takes place in the chloroplasts of plant vegetative tissues or in the plastids of non-photosynthetic ones. The metabolic pathways driving FA synthesis have been extensively studied in plants (for a review see Li-Beisson et al., 2013). Briefly, the pyruvate dehydrogenase complex (PDC) generates acetyl-CoA, the building block for FA production. FA biosynthesis begins with the formation of malonyl-CoA from acetyl-CoA driven by the heteromeric acetyl-CoA carboxylase (ACC). The malonyl group of malonyl-CoA is then transferred to an acyl-carrier protein (ACP), forming malonyl-ACP. Acyl chains are produced by the fatty acid synthase (FAS) complex, which uses acetyl-CoA as a starting unit, while malonyl-ACP provide the two carbon units required for chain elongation. Acyl chains are ultimately desaturated and/or hydrolyzed by stearoyl-ACP desaturase (SAD) and acyl-ACP thioesterases (FAT), respectively, releasing the free FA.

The use of DNA microarrays and transcriptome analyses has provided detailed information regarding the expression of genes involved in plant metabolic processes like FA biosynthesis (Schmid et al., 2005; Penouilh-Suzette et al., 2020). Hence, it appears that a number of the genes encoding core enzymes involved in FA synthesis are likely to be transcriptionally co-regulated (Mentzen et al., 2008). For instance, the induction of these genes is coordinated in embryonic tissues at the onset of seed maturation to ensure high levels of oil storage (Girke et al., 2000; Ruuska et al., 2002; Baud and Lepiniec, 2009). Transcription factors or proteins regulating mRNA turnover can control these changes in expression and indeed, genetic studies in Arabidopsis have revealed some of the factors that control seed oil biosynthesis. As such, several steps of FA synthesis are regulated by WRINKLED1 (WRI1: Ruuska et al., 2002; Baud et al., 2007, 2009; Maeo et al., 2009), a gene under the direct control of the transcription factor LEAFY COTYLEDON2 (LEC2: Baud et al., 2007), along with LEC1, FUSCA3 (FUS3) and ABA INSENSITIVE3 (ABI3), considered to be a master regulator of seed development (Braybrook and Harada, 2008; Suzuki and McCarty, 2008; North et al., 2010).

WRI1 encodes a transcription factor of the large APETALA2/ethylene-responsive element binding protein (AP2/EREBP) family (Cernac and Benning, 2004). Loss-of-function mutants have no obvious phenotype during vegetative development but they produce wrinkled, incompletely filled seeds, with an 80% reduction in seed oil content (Focks and Benning, 1998). In addition, these *wri1* mutants suffer a delay in embryo elongation and a modification of seed oil FA composition toward longer and more desaturated FAs (Baud et al., 2007). A combination of molecular and biochemical approaches identified possible WRI1-binding motifs in certain target genes (Baud et al., 2009; Maeo et al., 2009), and based on promoter sequence comparisons, the CNTNG(N)_7_CG motif was identified as the consensus WRI1 binding site and designated as the AW box (Maeo et al., 2009). WRI1 target sequences are found upstream of genes encoding for enzymes involved in glycolysis (sucrose synthase, pyruvate kinase and pyruvate dehydrogenase -PDH), subunits of ACC (biotin carboxyl carrier protein 2 - BCCP2- and its biotin attachment domain containing –BADC-homolog, carboxyltransferase -CT- and biotin carboxylase -BC), components of the FAS complex (malonyl-CoA:ACP malonyltransferase -MCMT, ketoacyl-ACP synthase -KAS, hydroxyacyl-ACP dehydrase -HAD, enoyl-ACP reductase -ENR- and ACPs), SAD, oleoyl-ACP thioesterase and genes involved in lipoic acid synthesis, a cofactor of PDH (Ruuska et al., 2002; Baud et al., 2007; Maeo et al., 2009; Pouvreau et al., 2011; To et al., 2012; Fukuda et al., 2013; Li et al., 2015; Liu et al., 2019; Kuczynski et al., 2020; Kazaz et al., 2020). More than 20 WRI1 target genes have been identified by comparing gene expression in wild type (WT) plants with that of *wri1* mutants and WRI1 overexpressing lines, and WRI1 promoter binding was confirmed in electrophoretic mobility shift assays (EMSA) and microscale thermophoresis experiments. The distance between the AW box and the translation initiation site (TIS) strongly influences the role of the AW box (Fukuda et al., 2013), and the majority of active AW sites lie less than 200 bp from the TIS in true WRI1 target genes (Maeo et al., 2009; Fukuda et al., 2013).

WRI1 orthologs have been identified in many plant species, including *Brassica napus* (Liu et al., 2010), *Zea mays* (Shen et al., 2010), *Elaeis guineensis* (Ma et al., 2013), *Brachypodium distachyon* (Yang et al., 2015), *Glycine max* (Chen et al., 2020) and *Oryza sativa* (Mano et al., 2019). The features of the *WRI1* gene, its transcriptional regulation and post-translational protein modifications have been studied widely in Arabidopsis, and in other dicotyledonous and monocotyledonous plants (Fei et al., 2020; Kong et al., 2020; Miray et al., 2021). Sunflower (*Helianthus annuus* L.) is an important oilseed crop cultivated worldwide. Like many other oilseed crops, several attempts to modify its oil composition have been made, for example, the development of new sunflower cultivars from mutagenized seeds with increased saturated FA levels (Zambelli et al., 2015). The biochemical pathways leading to oil synthesis and accumulation in sunflower seeds have also been studied widely (Salas et al., 2014; Venegas-Calerón et al., 2015). Analytical, biochemical, molecular and spatio-temporal expression studies (Martins-Noguerol et al., 2019 and 2020; Aznar-Moreno et al. 2016a, 2016b, 2018 and 2020; González-Thuillier et al., 2015, 2016 and 2021; González-Mellado et al., 2010, 2019) have enabled FA and TAG synthesis to be modelled in sunflower seeds, even though the regulation of these processes is less well characterized in sunflower. Recently, H3K4me3 epigenetic modifications were defined and related to the expression of FA-related genes, and of the VIV1 (homologous to Arabidopsis ABI3) and FUS3 transcription factors in developing sunflower seeds (Moreno-Pérez et al., 2021).

In this study we identified the sunflower *WRI1* gene and characterized its activity. We studied its role as a co-regulator of sunflower genes involved in plastidial FA synthesis, identifying the genes it recognizes and that are bound by this transcription factor. Interestingly, WRI1 mainly regulates very early steps of plastidial FA synthesis, particularly influencing the PDC subunits and the ACC complex, as well as acting through key FAS genes, driving the synthesis of oleic acid via *FATA1* gene regulation and establishing a push-pull strategy. We also analyzed the sequence of the functional sunflower AW box motif to determine if there is any base bias at the *N* positions of the sunflower AW box that drives active transcriptional regulation. As a result, we conclude that the sunflower AW box is sequence-sensitive at these non-conservative *N* positions.

## MATERIALS AND METHODS

### Plant material and growth conditions

Sunflower seeds from the common CAS-6 sunflower line (RHA-274 genetic background) were germinated in wet perlite at 25 °C and then moved to a germination chamber for 2 weeks. Subsequently, the seedlings were transferred to growth chambers and grown on 25 °C/15 °C (day/night) cycles in bags endowed with fertilizer. They were grown on a 16 h photoperiod with a photon flux density of 250 *μ*mol m^-2^ s^-1^.

Genomic DNA from mature sunflower leaves was extracted according to the modified cetyl-trimethyl ammonium bromide (CTAB) method (Zeinalzadehtabrizi et al., 2015). Total RNA was extracted from 21 days-after-flowering (DAF) sunflower seeds using the Spectrum Plant Total RNA Kit (Sigma-Aldrich, St. Louis, MO, USA), according to the manufacturer’s instructions. The total RNA obtained (1 *μ*g) was used to synthesize cDNA using the Ready-To-Go T-Primed First Strand Kit (Amersham Bioscience, Roosendaal, The Netherlands).

### HaWRI1-DNA binding domain cloning, expression in Escherichia coli and purification

The coding sequence of WRINKLED1 from the CAS-6 line (*HaWRI1*) was amplified by PCR from a cDNA pool of 21 DAF seeds using the HaWRI1-F and HaWRI1-R primer pair (Supplementary Table 1). The cDNA obtained was cloned into the pMBL-T vector (CANVAX Biotech, Córdoba, Spain), and its sequence was verified by DNA sequencing and deposited at GenBank (Acc. Number JX424422.1).

A truncated version of *HaWRI1* encoding the WRI1-DNA binding domain (*Ha*WRI1_DBD, amino acids 51 to 229), was cloned into the *pET_trx1a* expression vector (Baud et al., 2009) to obtain a 6-His-TRX-WRI1_DBD fusion protein. *Ha*WRI1_DBD was amplified with the Phusion High-Fidelity DNA polymerase (Thermo Fisher Scientific, Waltham, MA, USA) using the previously cloned full length *HaWRI1* cDNA as a template, and the HaWRI1-DBD-F and HaWRI1-DBD-R primers (Supplementary Table 1). The PCR product was digested and cloned into the *pET-trx1a* vector as a *Nco*I-*Xho*I fragment, removing the GFP coding sequence from this vector. The empty *pET-trx1a* vector provides the 6-His-TRX-GFP construct used in this work. The 6-His-TRX construct was obtained after digesting the *pET-trx1a* plasmid with the *Nco*I and *Xho*I restriction endonucleases (New England Biolabs, Hitchin, UK), removing the cDNA insert corresponding to the GFP coding sequence. After blunting the plasmid ends with Klenow enzyme (New England Biolabs, Hitchin, UK), the plasmid was religated with the T4 DNA Ligase (Thermo Fisher Scientific, Waltham, MA, USA). All the plasmids used here were verified by DNA sequencing.

Truncated protein expression and purification was carried out as described previously (Baud et al., 2009) with minor modifications. BL21(DE3) lacIq *E. coli* containing the *Ha*WRI1_DBD construct were grown at 37 °C to an OD_600_ of 0.6 in Luria Bertani (LB) medium containing NaCl (0.5 M) and protein expression was induced with 0.4 mM isopropyl-b-D-thiogalactoside (IPTG), allowing the bacteria to grow for an additional 20 h at 17 °C. The cells from a 1 liter culture were then harvested by centrifugation, washed with PBS (137 mM NaCl, 2.7 mM KCl, 10 mM Na_2_HPO_4_ and 1.8 mM KH_2_PO_4_ – phosphate buffered saline) and then sonicated in 8 ml lysis buffer (250 mM NaCl, 20 mM Tris–HCl [pH 8.0], 5 mM imidazole, 5% glycerol, 1 mM PMSF, 1/10 protein inhibitor cocktail: Sigma, St. Louis, MO, USA). After centrifugation for 40 min at 21,000 x*g*, the clear supernatant was incubated for 4 h with 1.5 ml Ni-NTA agarose (Qiagen, Hilden, Germany) previously washed with 6 ml of distilled H_2_O and equilibrated with 6 ml of binding buffer (150 mM NaCl, 20 mM Tris-HCl [pH 8.0], 2 mM MgCl_2_, 0.25 mM EDTA, 0.02% Nonidet P-40, 20% glycerol and 60 mM imidazole). The His-tagged recombinant protein-bound Ni-NTA agarose was transferred to an empty polypropylene column (Restek Corporation, Bellefonte, PA, USA) for gravity flow chromatographic purification, washed with 15 ml of washing buffer (binding buffer plus 60 mM imidazole) and the bound protein was eluted with 5 ml of elution buffer (binding buffer plus 300 mM imidazole). The fractions containing the protein were pooled and the imidazole removed on a PD-10 (sephadex G-25) column (GE Healthcare, Chicago, IL, USA), and the protein was concentrated in a centrifugal concentrator (Amicon MWCO 3 kDa: Merck, Darmstadt, Germany) and stored at −20 °C in dialysis buffer (150 mM NaCl, 20 mM Tris-HCl [pH 8.0], 2 mM MgCl_2_, 0.25 mM EDTA, 0.02% Nonidet P-40 and 20% glycerol). The purity of the recombinant protein was evaluated by SDS-PAGE and Coomassie staining (ChemiDoc Imaging System; BioRad, Hercules, CA, USA), and the concentration of the purified recombinant protein was determined with a Bio-Rad protein assay kit using bovine serum albumin (BSA) as a standard. Recombinant 6-His-TRX and 6-His-TRX-GFP proteins (used as a negative EMSA binding control), were obtained by the same procedure but starting from *E. coli* cells containing the 6-His-TRX construct or empty *pET-trx1a* vector, respectively.

### Promoter cloning, DNA fragment amplification and purification, and site-directed mutagenesis by overlap extension PCR

All the oligonucleotide sequences used in this work are shown in Supplementary Table 1 and were designed with the help of applications available on line: *primer3 4.0* (https://bioinfo.ut.ee/primer3-0.4.0/) and *OligoAnalyzer Tools* (https://eu.idtdna.com/pages/tools/oligoanalyzer). Oligonucleotide synthesis and DNA sequencing was performed by Eurofins Genomics (https://eurofinsgenomics.eu).

The promoter regions of the sunflower *FATA1*, *FATB1*, *SAD6*, *SAD17, KAR1* and *KAR2* genes (GenBank Acc. Numbers AY078350, AF036565, U91339.1, U91340, HM021135 and HM021136, respectively) were amplified by PCR from sunflower CAS-6 genomic DNA using the Phusion High-Fidelity DNA polymerase (Thermo Fisher Scientific, Waltham, MA, USA) and they were cloned into the pMBL-T vector (CANVAX Biotech, Córdoba, Spain). All the cloned promoter regions are located upstream of the TIS. The promoter length and the oligonucleotide pairs used for cloning are described in Supplementary Tables 2 (FAT and SAD genes) and 4 (KAR genes), and all the DNA clones obtained were verified by sequencing.

The promoter DNA fragments used here in the EMSAs are situated within -500 bp of the TIS of their corresponding gene, with some exceptions. Each DNA fragment was amplified by PCR from sunflower CAS-6 genomic DNA or the previously cloned promoter region as the template, using the Phusion High-Fidelity DNA polymerase (Thermo Fisher Scientific, Waltham, MA, USA) and the corresponding pair of oligonucleotides (see primer pairs used in Supplementary Tables 3 and 4, together with the product lengths and their location upstream of the TIS). PCR products were purified from agarose gels using the ISOLATE II PCR and Gel Kit (Bioline, Memphis, TN, USA), and quantified in a NanoDrop One C Spectrophotometer (Thermo Scientific, Waltham, MA, USA). All EMSA DNA PCR fragments were confirmed by sequencing.

Site-directed mutagenesis was performed by overlap extension PCR (Heckman and Pease, 2007) to introduce selected point mutations or DNA deletions into upstream sunflower DNA regions. The DNA fragment containing each wild type AW box sequence was used as the template for the first double PCR, and using the P1/P2 and P3/P4 oligonucleotide pairs, respectively, these containing the desired point mutation or deletion. The PCR products obtained were diluted 1/100, mixed and used as the template for extension and amplification in a second PCR with the P1/P4 oligonucleotide pair. The final PCR products consisted of the desired EMSA DNA fragment containing the mutated or deleted AW box. The correct introduction of the mutations was confirmed by sequencing (see Supplementary Table 5 for the mutations introduced, the EMSA DNA fragments generated and the mutagenic primer pairs used in each PCR -P1/P2, P3/P4, P1/P4).

### Agarose gel Electrophoretic Mobility Shift Assays (EMSAs)

Different amounts of the recombinant *Ha*WRI1_DBD fusion protein (6-His-TRX tagged, 0-640 ng range) were incubated for 30 min at room temperature with a single promoter DNA fragment (300 ng) in 20 μl of Binding Buffer (20 mM Tris-HCl [pH 8.0], 250 mM NaCl, 2 mM MgCl_2_, 1% glycerol, 1 mM DTT and 1 mg/ml BSA). Samples were then loaded onto a 1% agarose gel and resolved for 90 min at 4 °C at 100 V in 1× TAE buffer (40 mM Tris-HCl [pH 8.0], 20 mM acetic acid and 1 mM EDTA). The gel was soaked in 1× TAE buffer containing 0.5 μg/ml RedSafe (INtRON Biotechnology, Burlington, MA, USA) for 30 min and then visualized in a UV transilluminator.

### Acrylamide gel Electrophoretic Mobility Shift Assays (EMSAs)

The f1, f2 and f3 DNA fragments that correspond to the *FATA1* promoter region were designed to cover a 1 kb stretch relative to the ATG start codon in three overlapping fragments, each around 300-400 bp in length: f1, from −272 bp to +51 bp; f2, from –567 bp to –232 bp; and f3, from –918 bp to -534 bp. The *FATA1* promoter DNA fragment corresponding to f1 but without the AW box (*f1-ΔAWbox*, 299 bp) was generated by overlap extension PCR using the oligonucleotide pairs described in Supplementary Table 5 (see above). A 24 bp DNA fragment containing the 14 bp AW box sequence present in *FATA1* f1 was obtained by room temperature hybridization of the two complementary oligonucleotides pHaFATA1-AWbox-F and pHaFATA1-AWbox-R (Supplementary Table 1).

EMSA DNA fragments were labelled with digoxigenin (DIG) following the manufactureŕs instructions (DIG Gel Shift Kit 2^nd^ Generation; Roche Diagnostics, IN, USA) and 30 fmol of labelled DNA was incubated with different amounts of purified *Ha*WRI1_DBD recombinant protein (0-1000 ng) in 20 µl of Binding Buffer plus 0.05 µg/µl polydIdC. For competition assays, the unlabeled competitor was incubated with the protein briefly before adding the labelled DNA, and after adding the labelled DNA the reactions were incubated for 30 min at room temperature. The binding reactions were fractionated at 4 °C by electrophoresis at 80 V for 120 min on 6% native polyacrylamide gels (PAGE) in 0.5X TBE buffer (44.5 mM Tris-HCl [pH 8.0], 44.5 mM boric acid, 1 mM EDTA [pH 8.0]). Following electrophoretic separation, the oligonucleotide-protein complexes were transferred to a positively charged nylon membrane (Hybond-N+; Cytiva, Marlborough, MA, USA), and the DIG labelled DNA was visualized in an enzyme immunoassay using anti-Digoxigenin-AP and a chemiluminescent substrate, following the manufactureŕs instructions. The chemiluminescent signals generated were recorded on an imaging device (ChemiDoc Imaging System; BioRad, Hercules, CA, USA).

### Bioinformatics analysis

Protein sequences from known Arabidopsis genes involved in FA biosynthesis (http://aralip.plantbiology.msu.edu/pathways/fatty_acid_synthesis: Li-Beisson et al., 2013) were obtained from The Arabidopsis Information Resource, TAIR, (https://www.arabidopsis.org/: Berardini et al., 2015). These sequences were used to search for homologous genes in the sunflower genome, and to extract their promoter regions and transcriptomic data using the tools available in RELEASE 2.0 (v2020) of the Sunflower genome portal (Heliagene; INRA Sunflower Bioinformatic Resources, https://www.heliagene.org/: Badouin et al., 2017). The *in silico* localization of these proteins was predicted using DeepLoc (http://www.cbs.dtu.dk/services/DeepLoc/: Almagro-Armenteros et al., 2017) and Localizer (http://localizer.csiro.au/: Sperschneider et al., 2017) tools.

The DNA promoter regions were scanned to identify and localize AW box sequences (according to Maeo et al., 2009) using the PlantPAN 3.0 tool (http://plantpan.itps.ncku.edu.tw/: Chang et al., 2008). The Motif Alignment and Search Tool (MAST - http://meme-suite.org: Bailey and Gribskov, 1998) and WebLogo (https://weblogo.berkeley.edu/logo.cgi: Crooks et al., 2004) were used to obtained the consensus motif for the sunflower AW box.

## RESULTS

### The Sunflower genome contains one single WRI1 gene

The *A. thaliana* WRI1 protein (430 amino acids), encoded by the *At3g54320.1* gene (GenBank Acc. Number Q6X5Y6.1), was used as the query to search for *H. annuus* homologs in a public database (Heliagene) using the BLASTP algorithm. One particular protein sequence was detected with both the lowest E-value (2.63 x e^-109^) and the longest alignment (382 amino acids), corresponding to *HanXRQr2Chr14g0649121* gene located on chromosome 14. All the other homologs found had higher E-values, even for shorter alignments (174 amino acids or less). The percentage identity/similarity between AtWRI1 and the first four homologous sunflower proteins identified confirmed the *HanXRQr2Chr14g0649121* gene to be the sole homolog of *At*WRI1, with 50% identity and 64% similarity as opposed to less than 37% identity and 47% similarity for the other three genes detected. In addition, only *HanXRQr2Chr14g0649121* was expressed strongly in seeds, while the other three genes were preferentially expressed in leaves (Heliagene). Accordingly, we considered the *HanXRQr2Chr14g0649121* gene to be the *H. annuus* L. *WRINKLED1* gene and we refer to it hereafter as *HaWRI1*.

This *HaWRI1* gene shared the exon-intron structure of the *AtWRI1* gene (Figure 1A), which included the 9 bp exon 3 that encodes the VYL transcriptional activation motif (Ma et al., 2013), although a single base change (G/A) turned this motif into IYL in *Ha*WRI1. The highly conserved AP2 domain spanning all exons was also conserved in *HaWRI1*. The corresponding *HaWRI1* cDNA (1167 bp) from the CAS-6 sunflower line was cloned (see Methods) and its sequence was verified prior to depositing it in GenBank with the accession number JX424422.1. *Ha*WRI1 resulted in a protein 3 amino acids shorter (388) than that predicted in the Heliagene database, probably due to the distinct sunflower background (HA412-HO). Nevertheless, it included the main features of the WRI1 transcription factor (see Figure 1B for an alignment of the *At*WRI1 and *Ha*WRI1 proteins), the two AP2 domains (59-125 and 161-219 amino acids) and the two phosphorylation sites known to be involved in the protein stability (T64 and S160: Fei et al., 2020).

**Figure 1.**
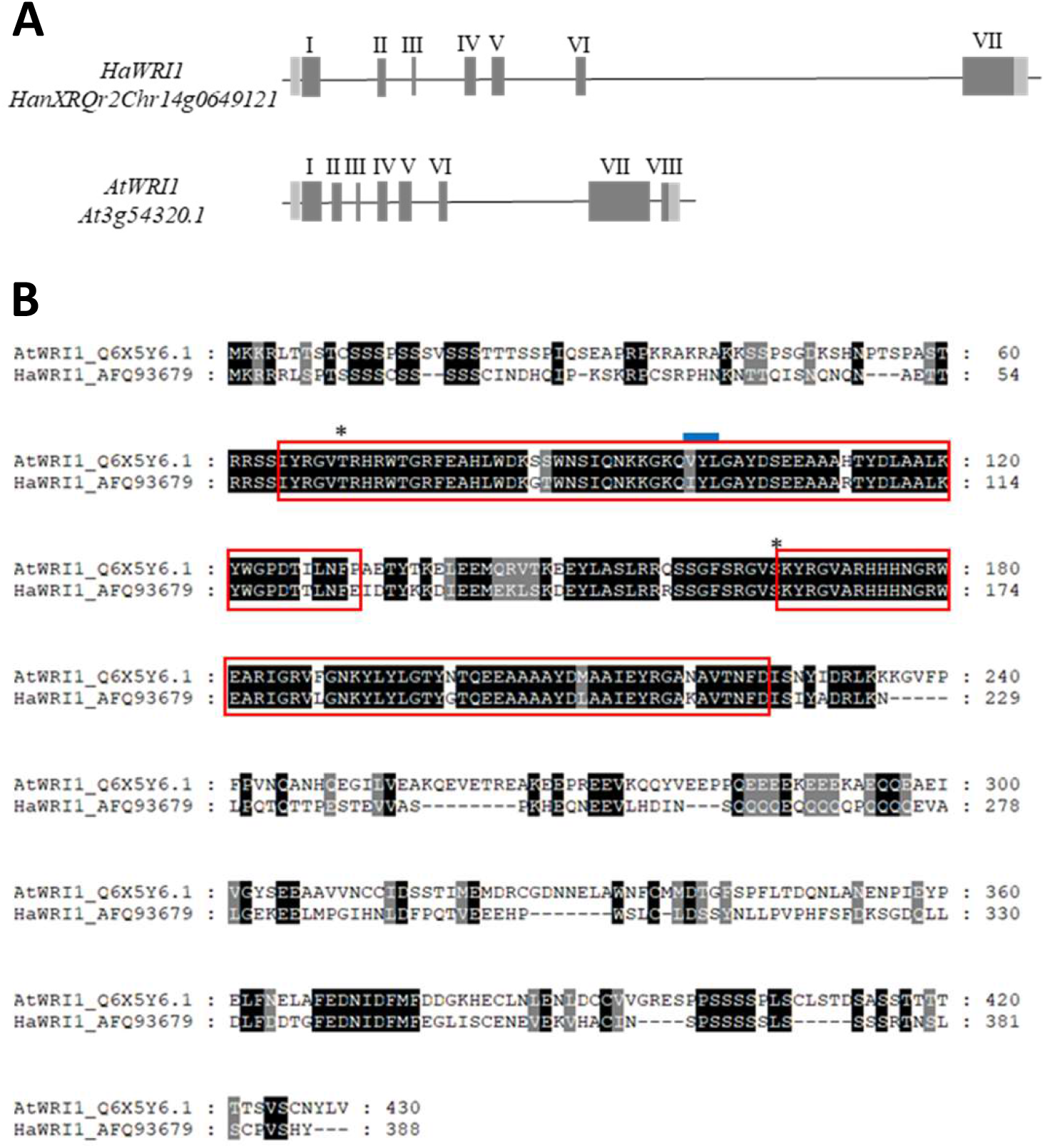
*Helianthus annuus* L. WRINKLED1. **A)** Arabidopsis (*AtWRI1*) and sunflower WRINKLED1 (*HaWRI1*) gene structures. **B)** *At*WRI1 and *Ha*WRI1 protein alignment. The AP2-EREBP DNA Binding Domain is highlighted by a red box and the asterisks mark the sites in *At*WRI1 phosphorylated by KIN10 (T70 and S166). The VYL domain encoded by 9 bp-exon 3 is marked with a blue line. The GenBank Acc. Numbers are shown on the left.

### Sunflower WRI1 binds to the AW box motif present in the plastid acyl-ACP thioesterase FATA1 gene

The primary function of WRI1 in seed oil deposition appears to be the positive regulation of genes that encode enzymes involved in late glycolysis and FA biosynthesis (Baud et al., 2009). The functionality of *Ha*WRI1 as a transcriptional activator was tested in EMSAs using DNA fragments containing the consensus AW box binding motif CNTNG(N)_7_CG (Maeo et al., 2009). To that end, we first cloned the DBD of *Ha*WRI1, fused it to thioredoxin (TRX) to improve its solubility and expressed it heterologously in *E. coli*, obtaining the 6-His-TRX-DBD recombinant protein (*Ha*WRI1_DBD). Moreover, we also expressed the 6-His-TRX and 6-His-TRX-GFP fusion proteins heterologously to use as negative binding controls in the EMSAs. All these recombinant proteins were purified by Ni-NTA affinity chromatography as described in the Methods (Supplementary Figure 1).

We next searched for candidate genes containing an AW box motif in their promoter region (as described by Maeo et al., 2009). We initially chose to work with FAT and SAD genes, not only due to their involvement in FA biosynthesis in seed plastids but mainly based on the *FATA* gene up-regulation in plants overexpressing WRI1 (Maeo et al., 2009; Li et al., 2015). Two sunflower FAT genes have been described in literature, *FATA1* and *FATB1*, with different substrate selectivity (Serrano-Vega et al. 2003 and 2005; Aznar-Moreno et al., 2016b), as well as two SAD genes, *SAD6* and *SAD17* (Hongtrakul et al., 1998; Serrano-Vega et al., 2003). We searched for the DNA promoter regions of each of these genes in the public sunflower database (Heliagene) using the BLASTN algorithm and the cDNA sequence of each gene as a query. Subsequently, primers were designed to amplify at least 1 kb of the promoter region for each gene, verifying the DNA fragments obtained by sequencing and used the PlantPAN 3.0 free software to search for the presence of the AW box (Supplementary Table 2). We found AW box motifs in the *SAD17*, *FATA1* and *FATB1* genes but not in *SAD6*. However, only *FATA1* presented an AW box motif in the upstream region close to the ATG codon, - 110/-97 bp from the TIS and in the 5’-untranslated region (5’-UTR), as described in Arabidopsis genes positively regulated by WRI1 (Maeo et al., 2009; Fukuda et al., 2013). All the AW boxes found, their sequences and locations from the TIS are detailed in Table 1, including a sequence similar to an AW box found in *SAD6* but with an extra base, together with the overlapping DNA fragments designed for EMSAs spanning the cloned promoter regions for each of the genes studied (Supplementary Table 3).

**Table 1.**
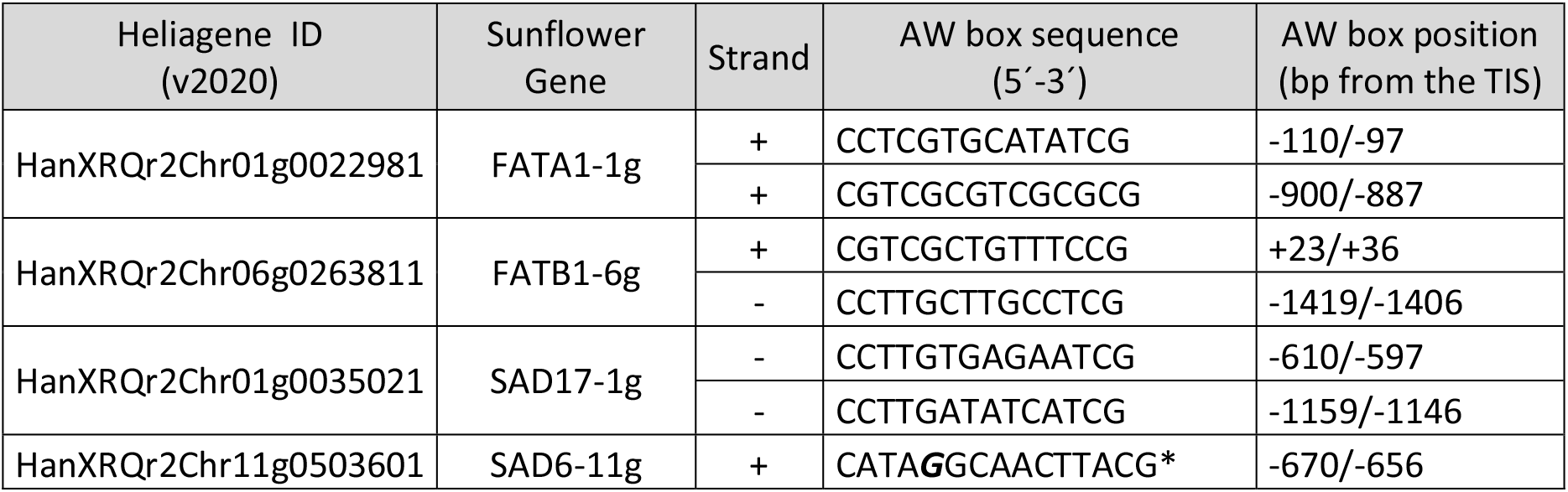
WRI1-binding motif (AW box: CNTNG(N)_7_CG) and their position relative to the ATG codon in the promoter regions of sunflower Acyl-ACP thioesterases (*FATA1*; *FATB1*) and Stearoyl-ACP desaturases (*SAD6*; *SAD17*). The asterisk marks a non-canonical sequence due to the appearance of an extra base (in bold) in the only motif found in *SAD6* similar to an AW box. The sunflower genes are named according to previous publications, followed by the chromosome number, as stated in this work for other sunflower genes: TIS, Translational Initiation Site (ATG codon).

EMSAs were performed to test the binding of the purified *Ha*WRI1_DBD to each overlapping DNA fragment of around 200-300 bp from the sunflower FAT and SAD promoter regions. This fragment size allowed us to resolve the potential shifted bands by electrophoresis in agarose gels. Binding of *Ha*WRI1_DBD to each DNA fragment from the promoter regions of the sunflower *FATA1* and *FATB1* genes (pHaFATs: Figure 2A), and the *SAD6* and *SAD17* genes (pHaSADs: Figure 2B), was analyzed and correlated to the locations of the AW box motifs in the overlapping DNA fragments (Figure 2C). As expected, the *FATA1* AW box at -110 bp from TIS included in fragment 1 (*pHaFATA1-f1*, 323 bp) bound to the *Ha*WRI1-DBD, migrating slower in the gels and increasing in intensity as the amount of protein assayed augmented, while the free DNA decreased concomitantly. Moreover, the shift of this fragment was not observed when *pHaFATA1-f1* was incubated with the GFP protein instead of WRI1. The mobility of none of the other DNA fragments tested was seen to alter in the gels whether or not they contained an AW box motif, neither was that of the *SAD6* sequences with an extra base (*SAD6-f4*) nor the specific binding control reactions with non-specific DNA, or when the GFP protein was used instead of WRI1.

**Figure 2.**
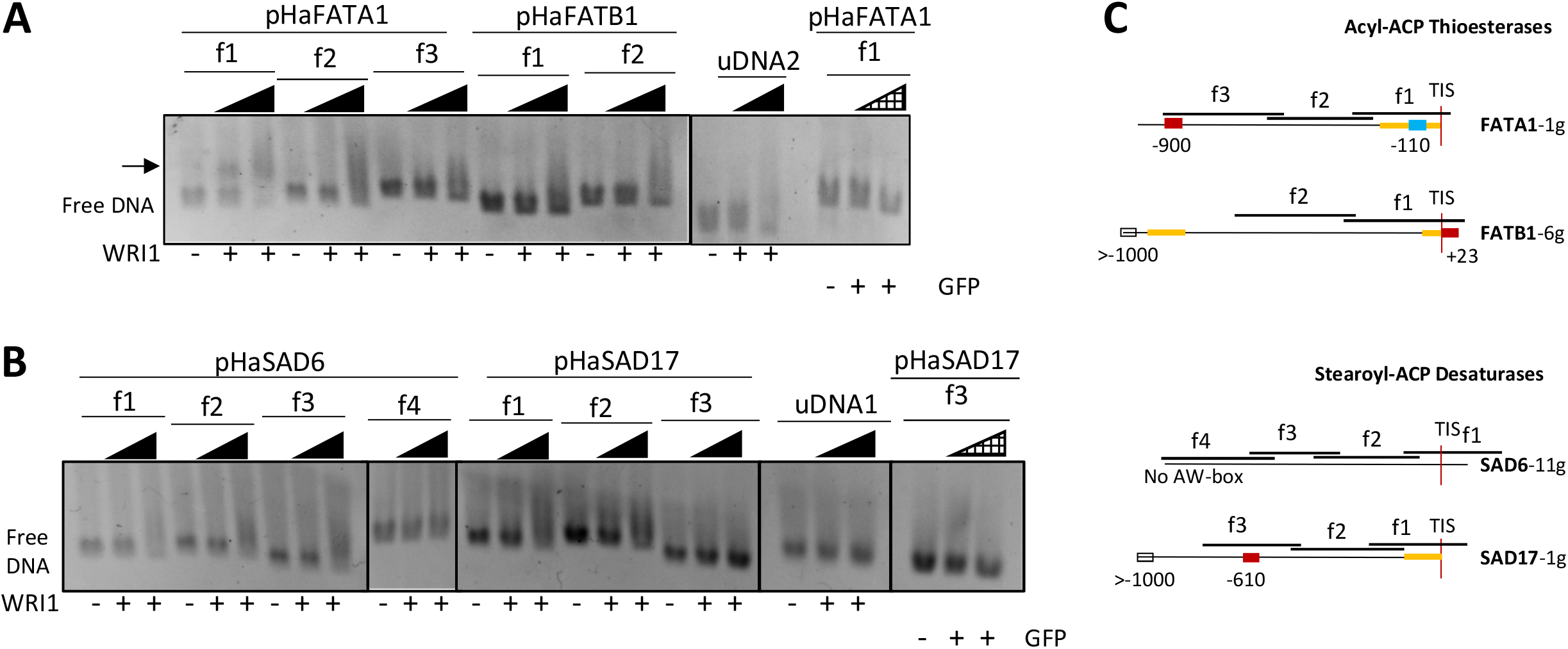
WRINKLED1 binding to the promoter regions (pHa) of sunflower (A) acyl-ACP-thioesterase (FAT) and (B) stearoyl-ACP desaturase (SAD) genes in agarose EMSAs. The arrow indicates the DNA shifted due to WRI1 binding. WRI1, 6-His-TRX-WRI1_DBD fusion protein (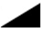 160-640 ng); GFP, 6-His-TRX-GFP fusion protein (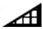 160-640 ng); uDNA1, non-specific DNA1 (*HacPGK2*); uDNA2, non-specific DNA2 (*HaCWI3*). C) Overlapping DNA fragments (f1 to f4, 200-300 ng) tested in each gene in agarose EMSA. The boxes indicate the AW box motifs and the numbers indicate the distance from the ATG codon (bp). Binding is indicated as negative (red boxes) or positive (blue boxes), while a colorless AW box means it was not tested. The 5’-UTR is in orange. The sunflower gene names are based on previous publications and they are followed by the number indicating the chromosome location. TIS, Translational Initiation Site (ATG).

To assess whether sunflower WRI1 bound specifically to the 14 bp AW box motif present in *pHaFATA1-f1* and not to any other sequence in the 323 bp f1 fragment, we performed EMSAs in acrylamide gels with digoxigenin labelled DNA incubated with the sunflower WRI1 DBD. The *Ha*WRI1-DBD bound specifically to double stranded DNA (dsDNA) fragments of 24 bp that contained the 14 bp f1 AW box motif from *pHaFATA1-f1* (the extra random bases were added to give a minimum 5 bp context at each end of the AW box motif, see methods). This shifted band was observed only in the presence of WRI1 and not when the DNA fragment was incubated with GFP or in competition assay with unlabeled dsDNA (Figure 3). Moreover, when *Ha*WRI1-DBD binding to the *pHaFATA1-f1* fragment lacking the 14 bp corresponding to the AW box (*f1-ΔAW-box*, 299 bp: Supplementary Table 5) was assessed in these EMSAs, no specific binding of this protein, or of the TRX and GFP fusion proteins, was observed (Supplementary Figure 2). Hence, sunflower WRI1 specifically bound to the AW box located at -110/-97 bp from the TIS in the 5’-UTR of *HaFATA1* gene.

**Figure 3.**
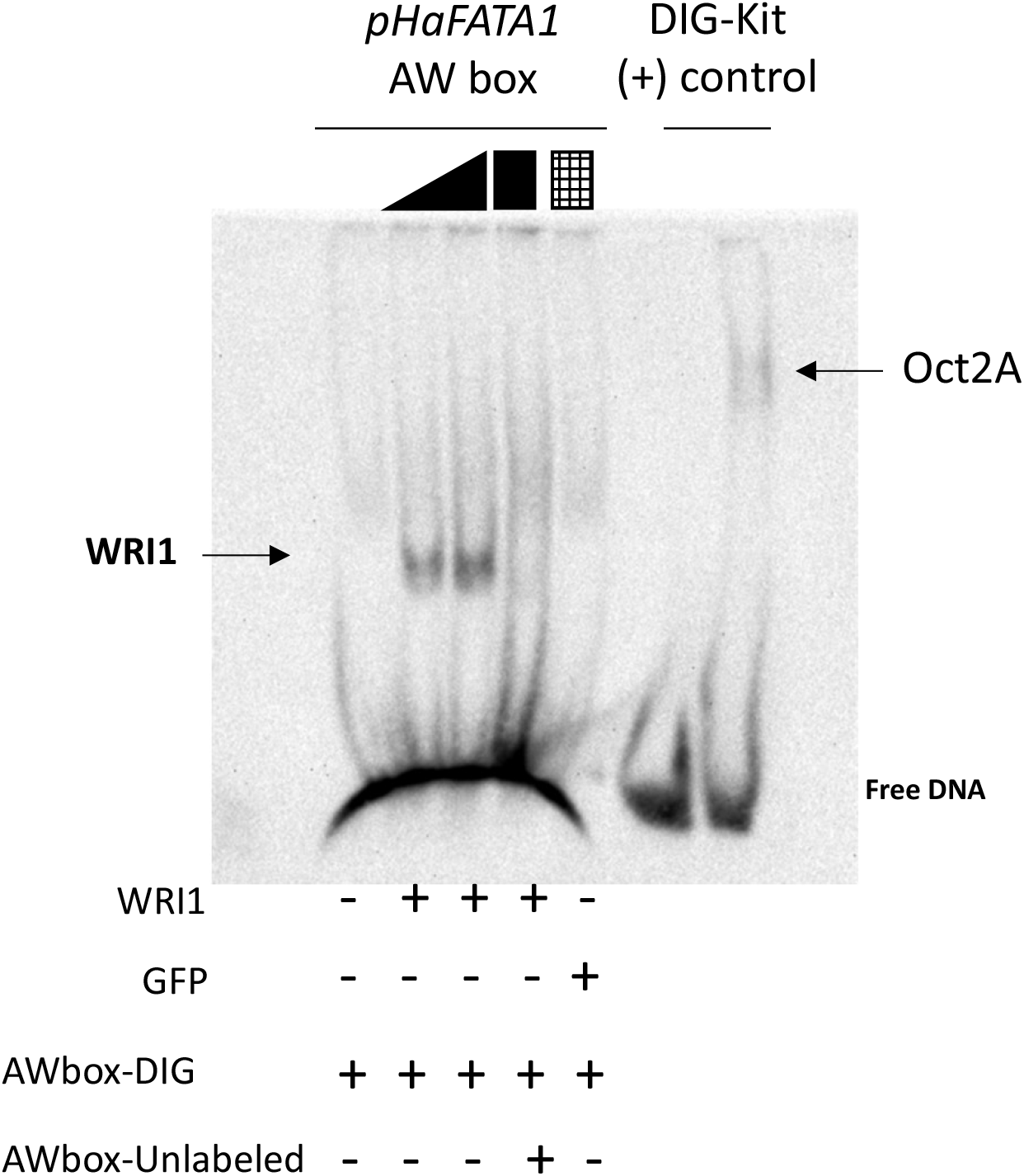
Binding of the HaWRI1_DBD to the AW box motif in the sunflower *FATA1* promoter region (*pHaFATA1*) as evident in Digoxigenin (DIG) labelled EMSA. The arrow shows WRI1 or Oct2A (positive-binding control) shifted DNA. The DIG-labeled or unlabeled (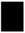 cold competitor reaction) dsDNA (24 bp, 30 fmol) contains the 14 bp AW box motif. WRI1, 6-His-TRX-WRI1_DBD fusion protein (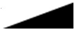 500-1000 ng); GFP, 6-His-TRX-GFP fusion protein (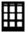 1000 ng); Oct2A, Octamer-binding factor 2A (75 ng).

### Sunflower WRI1 regulates plastid fatty acid synthesis mainly at early steps of the pathway

Having confirmed the functionality of *Ha*WRI1 by *in vitro* binding to the AW box motif located in *FATA1* upstream region, we investigated which other genes involved in sunflower intraplastidial FA biosynthesis might be regulated by WRI1 in a coordinated manner. In this regard, we were intrigued how WRI1 could discriminate the canonical AW box motifs also present in the DNA fragments *pHaFATA1-f3*, *pHaSAD17-f3* and *pHaFATB1-f1*, failing to bind to them in the *in vitro* binding assays (Figure 2C).

We used the BLASTN algorithm to search for the sunflower homologs of each gene involved in the plastidial FA biosynthetic pathway in the public sunflower database (Heliagene, recently updated in 2020) and using the corresponding Arabidopsis genes as queries (Table 2). The Arabidopsis gene sequences were obtained from The Arabidopsis Information Resource (TAIR) in accordance with the Arabidopsis Acyl-Lipid Metabolism Pathways (ARALIP). We assessed the existence of AW box motifs in the upstream regions of these genes using the PlantPAN 3.0 free software. We also employed two free online applications to predict the subcellular localization of all the genes (see Materials and Methods) and as expected, most of them were plastid genes. The genes with no AW box in their upstream region or with a subcellular localization other than plastid were ruled out for further study. The genes retained were referred to by their sunflower homologous gene name, the Heliagene ID, and for those with one or more AW boxes a shorter name was employed (the function acronym followed by a number with the chromosome location, e.g. *BCCP-16g*: Table 2). To be considered as a gene that could be regulated by WRI1 in the seed, we also analyzed the expression of each sunflower gene homolog in the seed based on the transcriptome data in the Heliagene database. According to this database (sunflower cv. *HA412-HO*), most genes were expressed in seeds and some of them strongly, such as the *β-PDH-16g*, *BCCP-9g*, *KASI-17g*, *KASIII-5g* and *KAR-17g* genes (Table 2). The seed expression data for all these genes was indirectly confirmed in the sunflower CAS-9 line as accessible chromatin in ChIP-seq experiments (Moreno-Pérez et al., 2021). Bibliographic references are given for those sunflower genes whose expression has already been described in CAS-6 sunflower seeds (Table 2). The expression of the other genes in seeds of the CAS-6 line, like the *FATB-9g* and genes of the sunflower ACC complex, was also confirmed (data not shown).

**Table 2.**
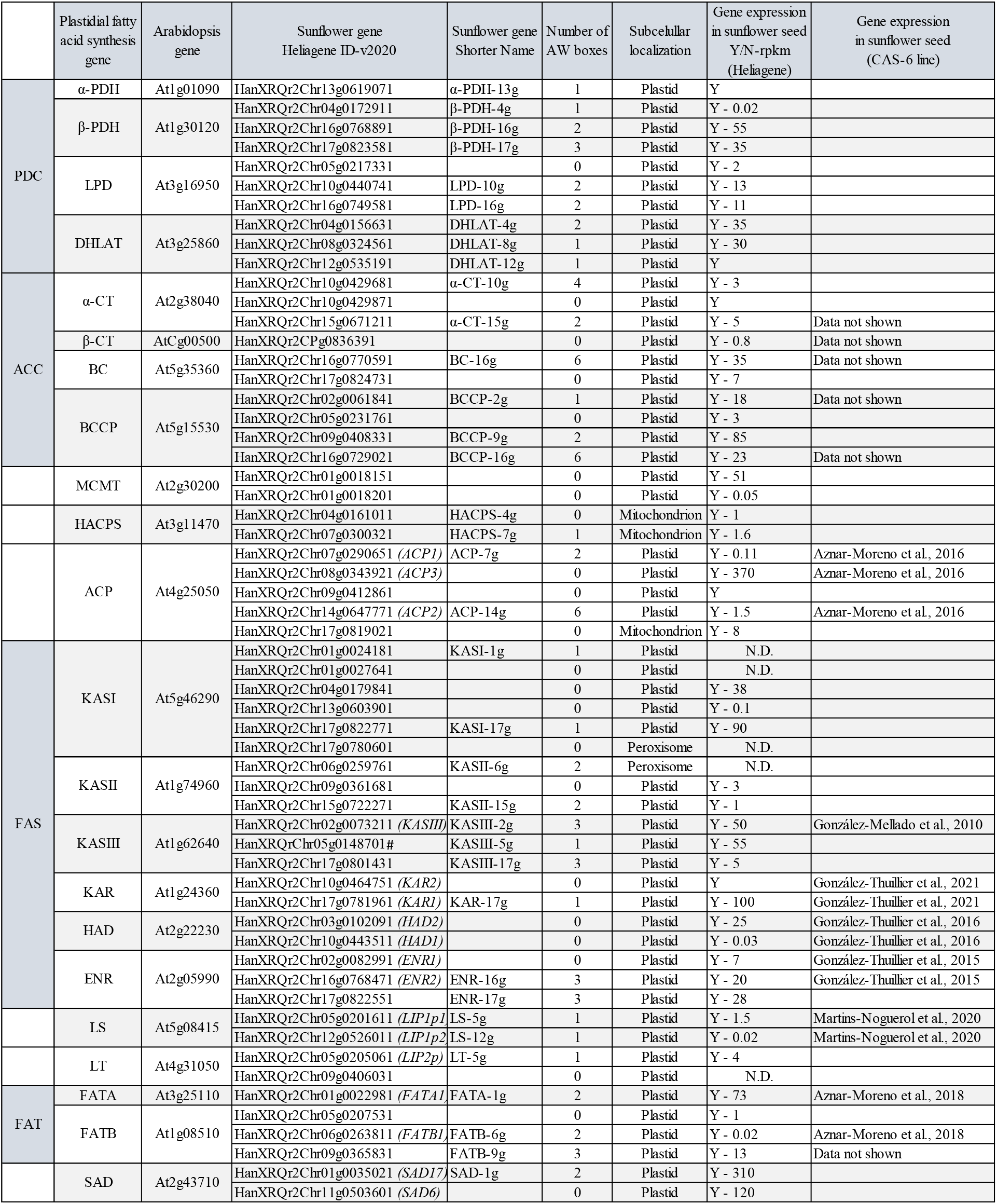
Sunflower genes involved in intraplastidial fatty acid biosynthesis. Arabidopsis genes were used as queries to search for sunflower homologs in a public database (Heliagene). The sunflower gene names are those taken from Heliagene ID (v2020) and with a shorter name used here according to the gene function acronym followed by the number of chromosome location: # Heliagene ID (v2018). Gene names already described in literature are indicated in brackets. The AW box motifs were found with the PlantPAN 3.0 tool. The subcellular localization was defined with the DeepLoc1.0 and Localizer free online applications. Gene expression in sunflower seeds is indicated according to the transcriptomic data from Heliagene-v2018 as YES (Y), indicating the transcript quantity as rpkm (reads per kilobase per million) when available (Heliagene-v2020). Bibliographic references are given when seed expression has previously been described in the CAS-6 line: N.D., No Data.

For all the genes expressed in seed plastids that contained one or more AW box, we situated the AW box motif in the upstream region relative to the TIS and determined if they fell within the 5’-UTR (Heliagene, see Table 3). We selected all the genes with AW boxes in 5’-UTR for further EMSAs, designing primers for PCR amplification and sequencing of each DNA fragment containing an AW box motif to use them in EMSAs (Supplementary Tables 4 and 5). As we wanted to test whether WRI1 could discriminate the consensus binding motif by its sequence alone, we included some AW box motifs found in upstream regions outside the 5’-UTR but close to the TIS (up to -500 bp) in the EMSA studies, and others found far from TIS (up to -2500 bp: see the genes, sequences and positions of all the AW boxes used in agarose gel EMSAs with sunflower WRI1 in Table 3). We were able to amplify and study all the selected AW box motifs, except for that in the *LT-5g* 5’-UTR (according to the Heliagene database), possibly because the sequence of the CAS-6 linés upstream region differs from that deposited in the database.

**Table 3.**
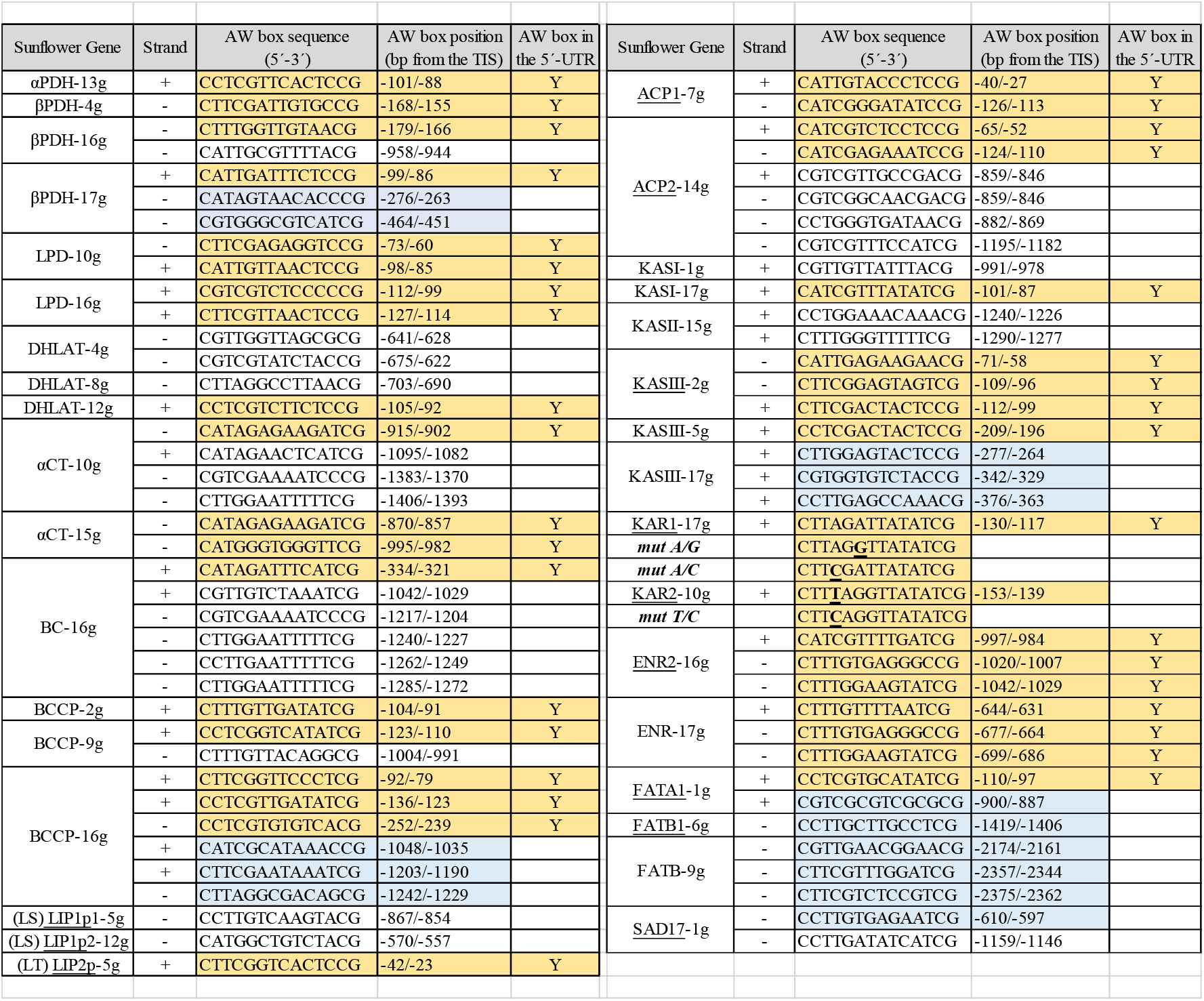
AW box motifs found in the promoter regions of sunflower genes probably involved in intraplastidial fatty acid biosynthesis. A search for the AW box motif was carried out using the PlantPAN 3.0 tool. The gene name is stated according to the functional acronym and chromosome location, and the position of the AW box motif is indicated as initial/end bp from the TIS and labeled with a YES (Y) when it lies within the 5’-UTR. Sunflower genes already described in literature are underlined. The sequences selected for EMSA are highlighted: Orange (located within 5’-UTR); Blue (located outside the 5’-UTR); # indicates the DNA sequence presents in *KAR2* promoter containing an extra base (underlined in bold) within the AW box motif. Point mutations in the KAŔs AW box motif are underlined in bold. Abbreviations: PDH, Pyruvate dehydrogenase; DHLAT, Dihydrolipoamide acetyl transferase; LPD, Dihydrolipoamide dehydrogenase; BC, Biotin carboxylase; BCCP, Biotin carboxyl carrier protein; CT, Carboxyltransferase; KAR, β-ketoacyl-ACP reductase; KAS, Ketoacyl-ACP synthase; ENR, Enoyl-ACP reductase; ACP, Acyl carrier protein; LS, Lipoate synthase; LT, Lipoyltransferase; FAT, acyl-ACP thioesterase; SAD, Stearoyl-ACP desaturase; TIS, Translational Initiation Site.

The results of the EMSAs with selected medium size DNA fragments (containing a single or multiple AW box motifs) were grouped according to the different enzymatic complexes or proteins that act in plastid FA synthesis. As such, PDC subunits encoded by the *α-PDH-13g*, *β-PDH-16g*, *β-PDH-17g*, *DHLAT-12g*, *LPD-10g* and *LPD-16g* genes contained an AW box motif in their 5’-UTR that bound the HaWRI1_DBD protein, and that produced a shift in EMSA agarose gels (Figure 4, panels A and B). The *α-PDH-13g*, *β-PDH-16g* and *DHLAT-12g* fragments each had a single AW box, whereas *β-PDH-17g* contained three boxes at -99 bp, -276 bp and -464 bp, although only that located within the 5’-UTR (-99 bp) gave a positive binding result. *LPD-10g* and *LPD-16g* each contained two AW boxes, both located in the 5’-UTR, yet while both the motifs in *LPD-10g* bound to *Ha*WRI1-DBD, only one motif in *LPD-16g* (at -127 bp) provoked a band shift. When analyzing the AW box motifs from the genes encoding the proteins of the ACC complex, the *BC-16g* and BCCP genes motifs were detected close to the ATG codon, whereas the α*-*CT genes contained motifs in their 5’-UTR but far from the TIS due to the presence of a 720 bp intron. Only *BCCP-9g* and *BCCP-16g* bound to *Ha*WRI1-DBD, producing a band shift in the assays (Figure 4, panels C and D). *BCCP-9g* had a single motif (at -123 bp) but *BCCP-16g* contained three motifs near the TIS (at -92 bp, - 136 bp and -252 bp), all in the 5’-UTR of the gene. When analyzed individually, the motifs at -136 bp and -252 bp were the only two DNA fragments that clearly underwent a shift in the presence of WRI1.

**Figure 4.**
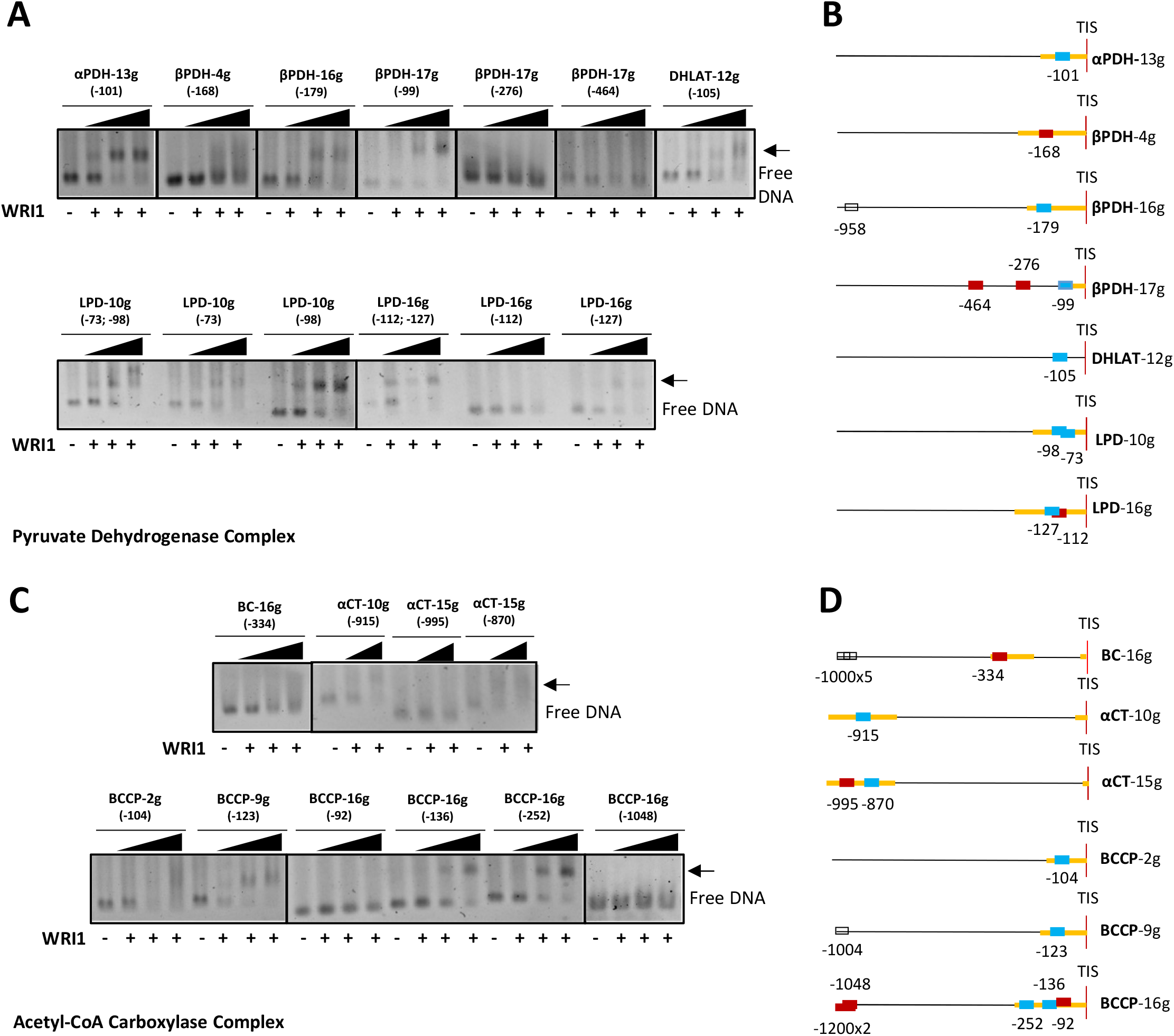
WRI1 binding in agarose EMSA to the sunflower promoter region of genes involved in plastidial fatty acid biosynthesis belonging to Pyruvate Dehydrogenase (A) and Acetyl-CoA Carboxylase (C) complexes. The arrow shows the DNA shift due to WRI1 binding: WRI1, 6-His-TRX-WRI1_DBD fusion protein (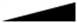 50-150-300 ng). **B** and **D**) Location of the AW box motifs relative to the ATG codon in each DNA promoter region analyzed by EMSA (300 ng). The boxes indicate AW box and the numbers indicate the location from the ATG (bp). Red boxes indicate negative binding and blue for positive binding, while a colorless AW box means it was not tested. The 5’- UTR is in orange. The sunflower gene names are based on previous publications and they are followed by the number indicating the chromosome location: TIS, Translational Initiation Site (ATG).

We also tested AW box motifs located far from the TIS in *BCCP-16g*, 3 motifs at >-1000 bp in the same EMSA, and as expected, no band shift was observed when these were incubated with *Ha*WRI1-DBD. The *BC-16g* AW box was located in the 5’-UTR and although no definitive band shift band there may have been some weak binding with the greatest amount of protein (smeared lane, Figure 4C). Indeed, the *α-CT-10g* and *α-CT-15g* genes contained an AW box in their 5’-UTR with an identical sequence (*αCT-10g* -915 bp and *αCT-15g* -870 bp), which underwent a shift in the EMSAs, whereas no binding was observed to the *αCT-15g* (-995 bp) motif. All these initial results confirmed that sunflower WRI1 could discriminate between canonical AW box motifs, specifically binding to just some of them in an *in vitro* assay and ruling out those located in the upstream region far from a TIS even though they might lie in the 5’-UTR. This phenomenon was also observed in the rest of the promoter regions analyzed (see below).

*Ha*WRI1_DBD binding to DNA fragments from regions upstream of the sunflower ACP genes was assessed, both *ACP-7g* (*ACP1*) and *ACP-14g* (*ACP2*) that contained two AW box motifs in their 5’-UTR (Figure 5, panels A and B). Only one AW box motif in each of these two genes produced a band shift, in both cases that located a little further away from the TIS (-126 bp in *ACP1* and -124 bp in *ACP2*). Regarding the AW box motifs in the upstream regions of genes belonging to the FAS complex (Figure 5, panels C and D), we first tested the motifs in selected KASIII genes (*KASIII-2g*, *KASIII-5g* and *KASIII-17g)* and *KASI-17g*. Of these, only the motif located in the 5’-UTR of *KASIII-5g* (-209 bp) produced a band shift when incubated with WRI1. The EMSA DNA fragment from *KASIII-2g* contained three AW boxes in the 5’-UTR, although they were too close together to be analyzed independently, and two of them overlapped and did not bind to WRI1, probably due to this overlap. The *KASI-17g* AW box motif produced a less clean but defined band shift. We also tested motifs located close to the ATG codon but in the 5’-UTR of the *KASIII-17g* upstream region and none bound to the *Ha*WRI1-DBD. Regarding the FAS complex, in addition to the KAS genes only the KAR and ENR genes contained WRI1-binding motifs. *ENR2-16g* and *ENR-17g* had three AW box motifs, two with an identical sequence (Table 2), and in EMSAs these gene fragments produced smears like *BC-16g* and *BCCP-2g*, suggesting some weak binding (Figure 4C). Hence, the sunflower genes targeted by WRI1 were mainly those involved in the early steps of the plastid pathways studied, including acetyl-CoA and malonyl-CoA synthesis, as all subunits of the PDC were encoded by genes bound by WRI1, along with the α-CT and BCCP genes of the ACC. Further down these pathways, a few key genes were bound by WRI1, such as ACP, KASIII, KASI and *FATA1*, affording the WRI1 transcript a co-regulatory role in sunflower seeds.

**Figure 5.**
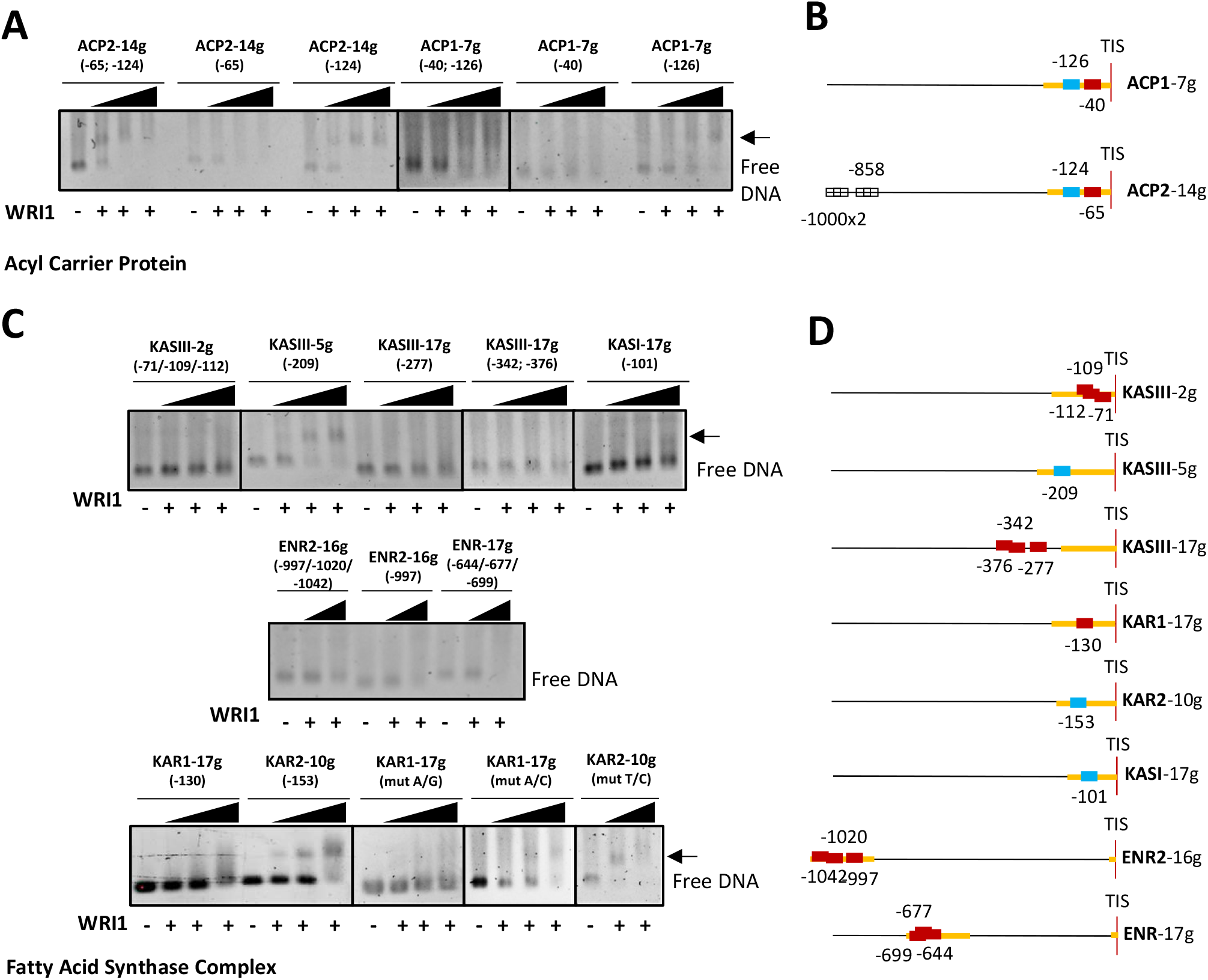
WRI1 binding in agarose EMSA to the promoter regions of sunflower genes involved in plastidial fatty acid biosynthesis, Acyl-Carrier Proteins (A) and those belonging to Fatty Acid Synthase Complex (C). The arrow shows DNA shifted by WRI1 binding: WRI1, 6-His-TRX-WRI1_DBD fusion protein (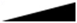 50-150-300 ng). **B** and **D**) Position of the AW box motif relative to ATG codon in each DNA promoter region analyzed by EMSA (300 ng). The boxes indicate the AW box motifs and the numbers indicate the position relative to the ATG (bp). The binding data are indicated as red for negative and blue for positive, while a colorless AW box means it was not tested. The 5’-UTR is in orange. The sunflower gene names are based on previous publications and they are followed by the number indicating the chromosome location: TIS, Translational Initiation Site (ATG).

### Sunflower WRI1 can bind to a non-canonical AW box motif

When considering the KAR genes, the *KAR-17g* (*KAR1*) gene contained a single AW box motif (Table 2) in its 5’-UTR at -130 bp, close to the TIS. Interestingly, the other sunflower KAR gene, *KAR-10g* (*KAR2*), did not have a canonical (CNTNG(N)_7_CG) AW box but rather, it has an almost identical sequence to that located in *KAR1* in its 5’-UTR (C<u>T</u>T<u>A</u>G<u>ATTATAT</u>CG: Table 3) that could be considered an “AW box with an extra “*N”* base (C<u>T</u>T*<u>T</u>*<u>A</u>G<u>GTTATAT</u>CG). To clarify the role of WRI1 in regulating these two isogenes that share 98.5% identity in their coding sequences and similar seed expression (González-Thuillier et al., 2021), we studied their promoter regions in EMSAs.

We amplified, cloned and sequenced DNA fragments corresponding to the upstream region of *KAR1* and *KAR2* using a common pair of primers due to the strong identity in their sequence, even in their 5’-UTR (Supplementary Table 4). The *KAR1* DNA fragment of 279 bp containing an AW box motif -130 bp from the TIS, while the 261 bp *KAR2* fragment contained a sequence similar to the AW box but with an extra base at -153 bp from the TIS. After incubating these two fragments with *Ha*WRI1-DBD we surprisingly observed a defined band shift in agarose gel EMSA. This *KAR2-10g* band become more intense as the amount of protein increased, whereas a smear developed when *KAR1-17g* DNA was incubated with the maximal amount of protein (Figure 5C, bottom panel). Specific binding control reactions with non-specific DNA and other purified proteins (GFP or TRX) produced negative results in EMSAs (Supplementary Figure 3). Hence, not only could *Ha*WRI1 bind to a non-canonical AW box motif *in vitro* but it also appeared to have a better affinity than to a canonical site in a similar context.

Given the strong identity between the two binding sequences assayed, differing only in one base (adenine at position N_3_ in *KAR1* and guanine in *KAR2*) and the presence of an extra base (thymine) in *KAR2* next to an adenine at position N_2_, we assessed whether the bases at N_3_ or N_2_ were crucial for WRI1 binding. We tested this using three sequences, two for the *KAR1* AW box and one for the *KAR2* motif (Table 3). For *KAR1*, the N_3_ base (adenine) was mutated to a guanine to imitate the *KAR2* AW box at that position (*mut A/G*), or the N_2_ base (adenine) was changed to a cytosine (*mut A/C*) as found in the *FATA1* AW box, also imitating an AW box bound by WRI1 *in vitro*. For *KAR2*, we mutated the extra base (thymine) adjacent to the N_2_ base to a cytosine (*mut T/C*) due to the presence of a cytosine at N_2_ in most AW boxes bound by sunflower WRI1. The binding of *Ha*WRI1-DBD to these mutated KAR AW boxes (Supplementary Table 5) was evaluated in agarose gel EMSAs (Figure 5C, bottom panel). The *KAR1 mut A/G* did not undergo a shift with WRI1 despite that mutation producing the same canonical sequence as that in *KAR2* except for the extra base, suggesting that the presence of the extra base in *KAR2* is responsible for the binding of WRI1, and that an adenine at the N_2_ position was not sufficient and the extra base was necessary for binding. However, *KAR1 mut A/C* did bind WRI1, producing a weak but visibly shifted band, reflecting the significant role of a cytosine at position N_2_ as expected. No difference was observed in the *KAR2 mut T/C* AW box compared to the WT and thus, the presence of an extra base next to the N_2_ adenine seemed to be the only requirement for WRI1 binding.

### The sunflower AW box motif that regulates transcripts involved in FA synthesis shows base bias in non-conservative motifs

We aligned the 52 canonical AW box motifs analyzed here in EMSAs classifying them according to their binding by sunflower WRI1 (Figure 6A). To identify the features enabling specific boxes to be recognized by and to bind to sunflower WRI1, we generated a position frequency matrix (PFM) and a position probability matrix (PPM) with the help of the MAST software available in the MEME suite using only the sequences that clearly bound WRI1 in EMSAs, 18 of the 52 analyzed. These matrices revealed missing bases and base bias at most non-conservative *N* positions of the sunflower AW box motif (Figure 6B). Thus, guanine was absent at the N_1_, N_2_, N_8_ and N_9_ positions, adenine was absent at N_4_ and cytosine was not detected at N_3_ and N_6_. These missing bases were considered as “forbidden” bases at those particular positions in the non-conservative part of the consensus AW box (as described by Maeo et al., 2009), suggesting that WRI1 recognizes and binds to the AW box motif in a sequence-sensitive manner in sunflower seeds. Regarding the base bias observed at most *N* positions, the prevalent bases at the N_2_ and N_6_ positions were cytosine and adenine, respectively. Further evidence of this sequence sensitivity came from the canonical AW box sequences not bound *in vitro* by sunflower WRI1 in the EMSAs, which all had one or more “forbidden” bases in their motif (Figure 6A, in red). When only one forbidden base was present it always involved position N_2_ (*KASIII-17g* -277 bp; *αCT-15g* -995 bp) or N_6_ (*BCCP-16g* -92bp; *KASIII-17g* -376 bp; or *ACP2-14g* -65 bp), highlighting the role and bias of both these positions. However, there were some exceptions in the absence of a “forbidden” base that could be explained by the presence/combination of one or more bases other than the critical N_2_ and N_6_ bias, such as *BC-16g* (-334 bp: N_2_=A; N_6_=T), *BCCP-2g* (-104 bp: N_2_=T), *KAR1-17g* (-130 bp: N_2_=A), *FATB-9g* (-2357 bp: N_6_=G) and *SAD17-1g* (-613bp: N_2_=T; N_6_=G). Moreover, when all the AW box motif sequences found in the sunflower genes that were not selected for EMSA due to their distance from the ATG codon (see Table 3) were examined, they mostly contained “forbidden” bases, and the three of these motifs that did not had one or more base at the N_2_/N_6_ position that was not considered among those within the WRI1 bias (*SAD17-1g*, -1159 bp; *KASII-15g*, -1290 bp; and *LS-5g*, -867 bp).

**Figure 6.**
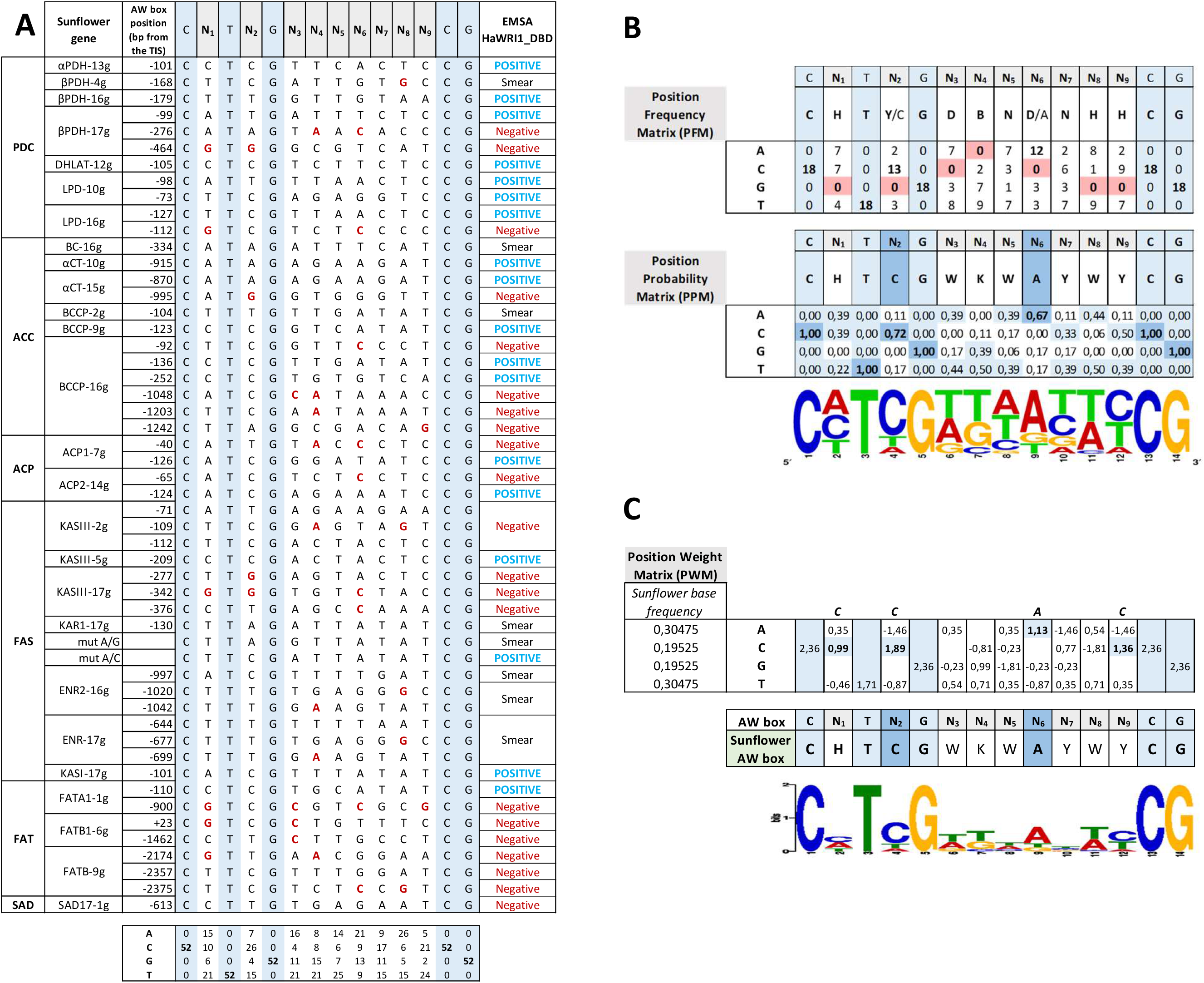
Sunflower AW box sequences present in genes involved in intraplastidial fatty acid synthesis. **A)** All the sunflower AW box motifs (CNTNG(N)_7_CG) analyzed by EMSA for binding to the sunflower WRINKLED1 DNA binding domain (*Ha*WRI1_DBD). The sunflower gene names, according to the functional acronym and chromosome location, and the AW box position relative to the ATG codon (TIS: Translational Initiation Site) are shown on the left. EMSA binding of WRI1 to each sequence is shown on the right and the “forbidden” bases are in red. **B)** The position frequency and probability matrices (PFM, PPM) for sunflower AW boxes that positively bind WRI1 reveal missing bases (red background) and the base bias (blue background) for most *N* positions of the canonical sequence. The motif derived from the PPM is shown as a WebLogo pattern. **C)** The position weight matrix (PWM) gives the sunflower base frequency and the active sunflower AW box motif is shown as a MEME pattern.

**Figure 7.**
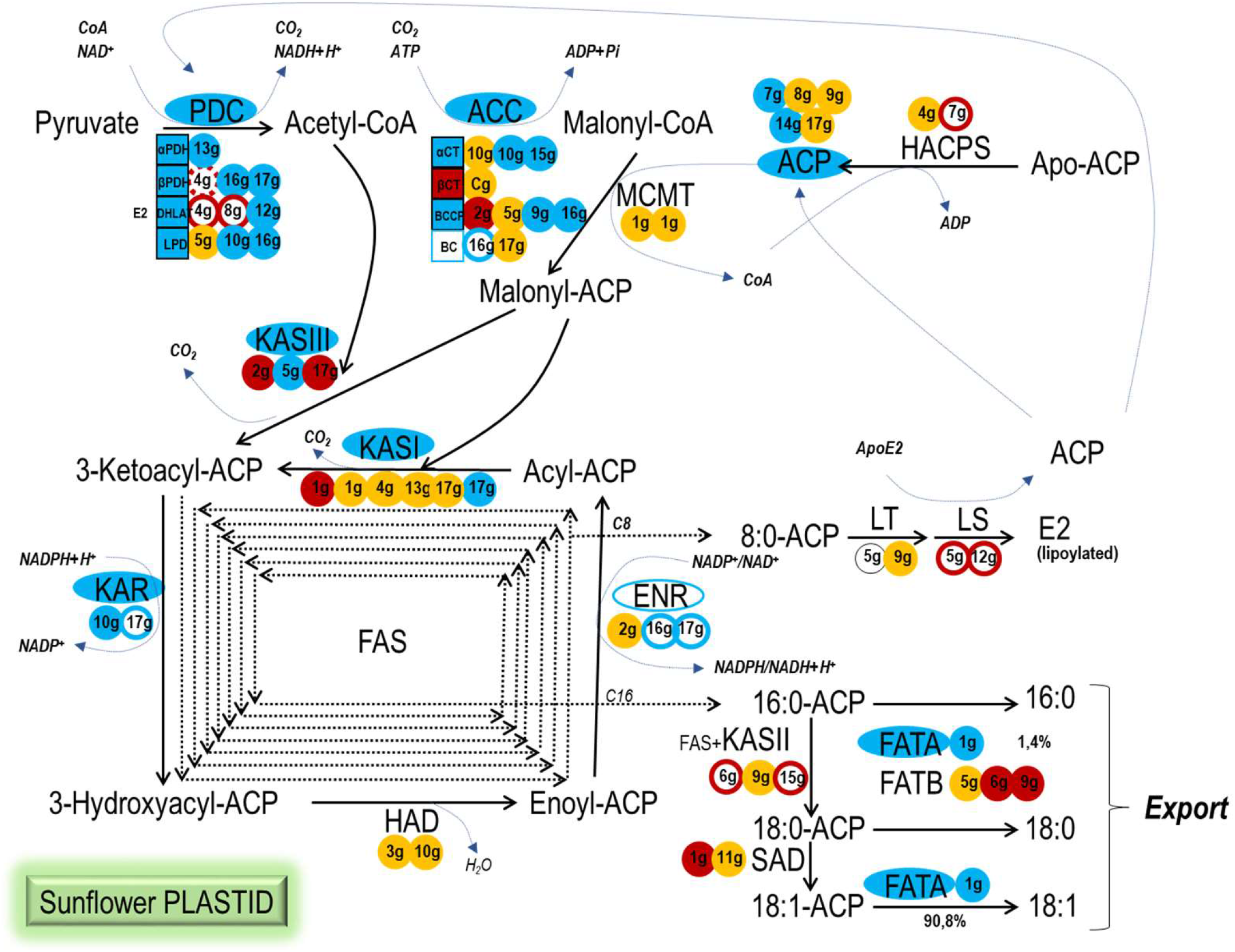
Sunflower WRI1 target genes in the seed plastid fatty acid biosynthetic pathway. WRI1 regulation of the sunflower gene isoforms studied here in EMSAs and the resulting active consensus sunflower AW box motif can be used to summarize the coordinated co-regulation of the pathway by WRI1: blue background, WRI1-EMSA positive; red background, WRI1-EMSA negative; blue outline, WRI1-EMSA smear, probably positive; dashed-red outline, WRI1-EMSA smear, probably negative; solid red outline, AW box motif outside the 5’-UTR and predicted not to be bound by WRI1; orange background, absence of a WRI1 AW box motif; black outline, AW box motif not present in the CAS-6 line according to the Heliagene database.

A new motif pattern was derived from the PPM (Figure 6B) and we wanted to fit this pattern to the sunflower genomic base frequency (Gill et al., 2014), for which we generated a position weight matrix (PWM) using the MAST software (Figure 6C). The observed base bias in the PWM corresponding to the 18 sunflower motifs produced a sunflower AW box motif that met the consensus motif (Maeo et al., 2009), yet it was fine-tuned to *<u>C</u>H<u>T</u>C<u>G</u>WKWAYWY<u>CG</u>*, highlighting the relevant role of the cytosine and adenine at the N_2_ and N_6_ positions, respectively.

## DISCUSSION

A high degree of cross-species conservation is a hallmark of master transcriptional regulators, as witnessed for WRI1 from *A. thaliana* (Cernac and Benning, 2004) and its orthologs in different plant species, both monocots and dicots. In most species there are mainly single WRI1 genes, except for *Z. mays* (Shen et al., 2010) and *C. sativa* (An et al., 2017) where two and three isoforms exist, respectively. Here we identified and studied the *WRI1* gene as a single copy gene in the genome from an important oil crop plant like *Helianthus annuus* (*HaWRI1)*. The *HaWRI1* gene and protein are consistent with those previously described for *AtWRI1* and its orthologs (Fei et al., 2020). However, it was notable that the third exon of *HaWRI1* encodes an IYL domain instead of the expected VYL due to a single (G/A) base change, also evident in some other WRI1s like as the tomato *Sl*WRI1 protein (GenBank XP004231231). The VYL domain is a highly conserved domain and site-directed mutagenesis of this motif in *At*WRI1 failed to restore the oil content in the *wri1* mutant (Ma et al., 2013). However, WRI1 orthologs identified in *R. communis* (Tajima et al., 2013) and *O. sativa* (Mano et al., 2019) encode proteins lacking a VYL motif, yet oil biosynthesis *in planta* was still triggered by activating the expression of WRI1 target genes (Ji et al., 2018; Mano et al., 2019). IYL is thought to be produced through a G to A substitution in the first codon of the motif for “V”, as occurs in *Ha*WRI1. As such, the first amino acid “V” seems to be less important in this VYL motif and can be substituted (Ma et al., 2013; Krizek, 2003).

Some WRI1s are expressed strongly in non-seed tissues, such as fruits (e.g. oil palm WRI1). However, numerous WRI1-like proteins are expressed strongly in developing seeds, closely correlated to *AtWRI1* expression. As expected for the sunflower, based on the sunflower transcriptome data (Heliagene), *HaWRI1* was most strongly expressed in seed tissues. When searching for WRI1 homologs in the sunflower database using *AtWRI1*, other putative WRI genes were found with higher expression levels in non-seed tissues (leaves, ligule), probably members of the sunflower WRI gene family whose activity remains to be studied. Up to four members of this family have been described in *A. thaliana* (*AtWRI1-4*: To et al., 2012), with *AtWRI2-4* expressed in floral organs and active in cutin biosynthesis, and there are three family members in *R. communis* (*RcWRI1-3*, Tajima et al., 2013), acting in membrane lipid synthesis in vegetative organs.

WRI1 is described as a master regulator of genes involved in late glycolysis and FA biosynthesis in *Arabidopsis* seeds (Baud et al., 2009). The activity of *Ha*WRI1 as a transcriptional regulator of sunflower genes encoding enzymes involved in seed plastid FA biosynthesis was confirmed in EMSAs using DNA fragments containing the consensus AW box motif found in the upstream regions of the sunflower genes studied: CNTNG(N)_7_CG (Maeo et al., 2009). These EMSA results indicate that sunflower WRI1 mainly drives and coordinates the early steps of FA synthesis in seed plastids. All the subunits of the PDC complex are encoded by genes under the control of WRI1, as are the genes encoding for the αCT and BCCP subunits of the ACC complex. Later in the pathway, WRI1 regulated the transcription of two out of three ACP genes, and in the FAS complex, only the genes encoding KASIII, KASI and KAR activities clearly bound this transcription factor, unlike other FAS complex genes like HAD or KASII. Interestingly, WRI1 also regulated the *FATA1* gene but no *FATB* gene, driving the sunflower seed toward the synthesis and export of oleic acid, although no SAD gene was found to be controlled by WRI1. This regulatory framework for FA synthesis in the seed plastid driven by *Ha*WRI1 was consistent with that stated in the literature for Arabidopsis (Ruuska et al., 2002; Baud et al., 2007; Maeo et al., 2009; To et al., 2012; Fukuda et al., 2013; Kuczynski et al., 2020), as well as for *B. napus* (Li et al., 2015) and other crops like maize (Pouvreau et al., 2011). In Arabidopsis, a combination of microarrays, RT-PCR, yeast-two-hybrid screening, EMSA and thermophoretic analysis, together with *in vivo* GUS experiments, established WRI1 target genes that included all the sunflower genes revealed here in EMSAs. Some of these, like the KASI, ACP, *FATA* and *BCCP2* genes, were also described as candidate targets in *B. napus* or *Z. mays*. However, our confirmed candidates in sunflower did not include all the targets described in Arabidopsis that are involved in plastid FA biosynthesis, such as MCMT and HAD, as well as those involved in lipoic acid synthesis (LS, LT) and SAD genes (Kuczynski et al., 2020; Kazaz et al., 2020; To et al., 2012; Pouvreau et al., 2011; Baud et al., 2007). MCMT is also described as a putative WRI1 target gene in *B. napus* because it is upregulated in plants overexpressing *Bn*WRI1, although neither this nor the HAD gene had an AW box in their upstream regions in the sunflower genome.

Regarding sunflower LS, LT and SAD genes, some of their sunflower isoforms had AW box motifs in their upstream regions but outside of the 5’-UTR and far from the TIS, except for a putative motif at -42 bp for the *LT-5g* gene that was not detected in the genome of the sunflower CAS-6 line. The distance between the AW box and the TIS strongly influences the function of the AW box (Maeo et al., 2009; Fukuda et al., 2013), and the majority of AW sites lie within 200 bp of the TIS in WRI1 target genes. Accordingly, when we tested the sunflower AW box motifs far from the TIS, including those in the SAD genes, they all failed to bind *Ha*WRI1. Although multiple and diverse techniques have been used to identify the WRI1 target genes, the presence or absence of a specific target may reflect the technical sensitivity of these tests. In addition, we cannot rule out that the minor differences in WRI1 in terms of the regulation of FA synthesis in seed plastids may be specific to the plant species. Hence, the HAD, LS and SAD genes in another crop species like maize are not *Zm*WRI1a target genes (Pouvreau et al., 2011), like we describe for sunflower.

Both the BC and ENR genes are described as *At*WRI1 targets in binding assays of the AW box motif (Maeo et al., 2009; Kuczynski et al., 2020), yet it was unclear if they are *Ha*WRI1 target genes based on the EMSA results alone, as a smear as opposed to a clear band shift was evident for both. However, the sequence analysis of their AW box motifs suggests that they may be regulated by WRI1 as they present at least one AW box in their 5’-UTR that was consistent with our sunflower AW box consensus motif. Hence, the smeared lanes might be explained by less stable DNA binding and thus, we should not rule them out as potential sunflower WRI1 targets. The *in silico* presence of an AW box motif in the sunflower *KAR1* upstream region but not in *KAR2* suggests that *KAR1* gene but not *KAR2* is a putative WRI1 target in sunflower seeds (González-Thuillier et al., 2021). The corresponding gene in Arabidopsis (*30AR*, *At1g24360*) is also a putative WRI1 target due to its upregulation in WRI1-overexpressed plants (Maeo et al., 2009). We amplified and sequenced the upstream region of both sunflower genes, confirming the presence of the AW box motif in the *KAR1* upstream region but also, detecting the presence of an almost identical sequence in the corresponding *KAR2*, albeit an AW box with an extra base. Unexpectedly, the non-canonical *KAR2* AW box motif was clearly bound by *Ha*WRI1, as opposed to the *KAR1* canonical motif that produces a smear in the EMSA gel. Considering that the *KAR1* AW box conforms to the sunflower consensus motif, as occurred in the BC and ENR genes, and given the EMSA results for both genes, we propose that *KAR1* and *KAR2* are indeed *Ha*WRI1 targets with possible differences in their affinity to the transcriptional regulator. Moreover, *in vitro Ha*WRI1 binding to a non-canonical AW-box opens the possibility of finding new WRI1 target genes and/or the number of binding sites in a promoter region. The two binding sites involved in sunflower KAR genes had almost identical sequences, yet EMSA with specific mutations at *N* positions revealed that base bias at N_2_ plays an important role in the binding stability between these *cis* elements and the regulator. Moreover, the selective *in vitro* binding of sunflower WRI1 to only some of the AW boxes tested suggests that WRI1 could act in a sequence-sensitive manner relative to the bases at *N* positions within the AW box, in addition to the *in vivo* position-sensitive binding described previously (Fukuda et al., 2013).

The AW box motif CNTNG(N)_7_CG was first described as the *cis At*WRI1 binding element (Maeo et al., 2009), and it was later studied in other plants like *Z. mays* (Pouvreau et al., 2011) and *B. napus* (Li et al., 2015), confirming this motif as the consensus WRI1 binding motif in plants. Most previous studies focused on the presence/absence of this motif as a seal of the putative regulation of transcription by WRI1 when located close to the transcriptional or translational start site. In sunflower, we found this *cis* element in the upstream regions of seed plastid FA synthesis genes, yet only 18 of 52 were recognized and bound by *Ha*WRI1 in *in vitro* EMSAs. Most of the motifs not bound by WRI1 were located far from the TIS of the corresponding genes but others were close to a TIS or at least within the 5’-UTR. Hence, the motif’s location is not sufficient to determine if WRI1 will bind to a motif but probably, this also depends on the sequence itself, in which certain non-conservative positions play an important role. Our analysis of the sunflower AW box sequences recognized by *Ha*WRI1 *in vitro* revealed some forbidden bases at specific *N* positions, and a particularly relevant base bias for the N_2_ and N_6_ positions. From this analysis a consensus sunflower AW box motif could be drafted that was active in the regulation of FA biosynthetic gene transcription by WRI1 in seed plastids.

These active sunflower seed motifs were mainly found as single binding boxes in the gene’s promoter region and only two genes were found to have two active AW boxes in their upstream regions, *LPD-10g* and *BCCP-16g*. In Arabidopsis, the 5’-UTR of *BCCP2* contains two AW boxes and expression of the *BCCP2* gene is much more strongly induced by WRI1 relative to genes with a single AW box (Maeo et al., 2009). According to the transcriptomic data for the common sunflower (Heliagene), the *LPD-10g* and *BCCP-16g* genes were expressed strongly in seeds, with only a slightly higher number of transcripts in ligules and roots, respectively, yet at a similar level to other genes that have only a single AW box, *LPD-16g*, *BCCP-2g* and *BCCP-9g*. Thus, there may be other factors playing a role in regulating the transcription of genes involved in FA synthesis in sunflower seeds. Nevertheless, the active motifs identified did indicate that the PDC (LPD, β-PDH), ACC (BCCP) and KASIII gene families expressed strongly in sunflower seed contain active AW motifs in their promoter regions (genes that clearly shifted in EMSAs). By contrast, *LPD-5g* and *BCCP-5g* had no AW motifs and they were expressed weakly in seeds compared to other members of their families. The β-PDH and KASIII genes all contain AW boxes and are expressed more strongly in seeds when they contain an active motif, such as *β-PDH-16g* and *β-PDH-17g* as opposed to *β-PDH-4g*, and *KASIII-2g* and *KASIII-5g* as opposed to *KASIII-17g*. However, it does not appear to be a general rule as the ACP gene expressed most strongly in seeds was that with no AW motif in its promoter region, the *ACP-8g* (*ACP3*) gene. These results confirmed that while other regulatory factors might be active in these events, the use of WRI1 to regulate FA synthesis in sunflower seed plastids focuses mainly on early stages of the pathways involved, that of acetyl-CoA and malonyl-CoA synthesis by the PDC and ACC complexes, respectively, together with the first step in the FAS complex, 3-ketoacyl-ACP synthesis by KASIII. Moreover, the fact that the single gene coding for *Ha*FATA1 was the only thioesterase gene bound by *Ha*WRI1 in EMSA, and none of the FATB genes were retarded by this transcription factor (*FATB1* or *FATB-9g*), leads us to propose a push-pull strategy employed by sunflower WRI1 to regulate seed plastid FA synthesis, directing it toward the synthesis and export of oleic acid.

### Future Perspective

The specific characterization of WRI1 targets in sunflower seeds paves the way to redirect their FA synthesis by modifying/adapting gene promoters not regulated by this factor using gene editing techniques, such as CRISPR-CAS9. In addition, the recognition of non-canonical boxes by *Ha*WRI1 opens the door to new studies on the amino acid residues involved in such events, and/or the ability of WRI1s from other species to recognize these new targets with an extra nucleotide.

## Abbreviations

ACC: acetyl-CoA carboxylase complex
ACP: acyl carrier protein
BC: biotin carboxylase
BCCP: biotin carboxyl carrier protein
CT (α and β subunits): carboxyltransferase
DAF: days after flowering
DHLAT (E2): dihydrolipoamide acetyl transferase
dsDNA: double stranded DNA
EMSA: electrophoretic mobility shift assay
ENR: enoyl-ACP reductase
FA: fatty acid
FAS: fatty acid synthase complex
FAT: acyl- ACP-thioesterase
HACPS: holo-ACP-synthase
HAD: hydroxyacyl-ACP dehydrase
KAR: β-ketoacyl-ACP reductase
KAS: β-ketoacyl-ACP synthase
LPD (E3): dihydrolipoamide dehydrogenase
LS/LIP1: lipoate synthase
LT/LIP2: lipoyltransferase
MCMT/MAT: malonyl-CoA-ACP malonyl transferase
PDC: pyruvate dehydrogenase complex
PDH (E1, α and β subunits): pyruvate dehydrogenase
PFM: position frequency matrix
PPM: position probability matrix
PWM: position weight matrix
SAD: stearoyl-ACP desaturase
TAG: Triacylglycerol
TIS: translation initiation site
UTR: untranslated region
WRI1: Wrinkled1
WT: wild type

## Acknowledgements

We thank M.A. Troncoso-Ponce (Université de Technologie de Compiègne, France) for kindly providing the *pET-trx1a* vector and A. González-Callejas for skillful technical assistance. This research was funded by the Spanish AEI/FEDER (UE), Projects AGL2017-83449-R and PID2020-113134RBI00/AEI/10.13039/501100011033.

## Author contribution statement

Rosario Sánchez and Enrique Martínez-Force conceived the experimental design and wrote the manuscript, Rosario Sánchez performed the experiments, Irene González-Thuillier did the full-length cDNA cloning of *HaWRI1*, Monica Venegas-Calerón, Joaquín J. Salas and Rafael Garcés revised the manuscript. All authors read and approved the manuscript.

## Supplementary data

**Suppl. Fig. 1. SDS-PAGE followed by Coomassie staining to show the purified recombinant proteins used in EMSA.** *Ha*WRI1_DBD, sunflower 6-His-Thioredoxin-WRINKLED1_DNA Binding Domain (312 aa, 35 kDa); TRX, 6-His-Thioredoxin fusion protein (163 aa, 18 kDa); GFP, 6-His-Thioredoxin-Green Fluorescent Protein (773 aa, 41 kDa). MW, Molecular Weight; T, total fraction; SB, soluble fraction; P, purified protein.

**Suppl. Fig. 2. *Ha*WRI1_DBD binding to DNA fragment 1 of the sunflower *FATA1* promoter region (*pHaFATA1-f1*) containing an AW box motif and specific control binding in Digoxigenin (DIG) labelled EMSA.** The arrow shows the DNA shifted by WRI1 binding. DIG labeled or unlabeled (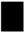 cold competitor reaction) dsDNA (30 fmol): f1 (323 bp) and f1-ΔAWbox (299 bp) with or without the 14 bp AW box motif, respectively. WRI1, 6-His-TRX-WRI1_DBD fusion protein (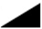 160-800 ng); GFP, 6-His-TRX-GFP fusion protein (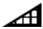 160-800 ng); TRX, 6-His-Thioredoxin protein (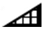 160-800 ng).

**Suppl. Fig. 3. *Ha*WRI1_DBD binding to the sunflower β-ketoacyl-ACP reductase (*KAR1* and *KAR2*) promoter regions and specific control binding reactions in agarose EMSA.** The arrow shows the DNA shifted by WRI1 binding, the DNA (300 ng) consisted of the corresponding promoter region containing the AW box motif studied, as indicated by its location from ATG codon: WRI1 (W), 6-His-TRX-WRI1_DBD fusion protein (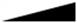 150-300-600 ng); TRX (T), 6-His-Thioredoxin protein (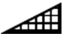 150-300-600 ng); GFP (G), 6-His-TRX-Green Fluorescent protein (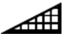 150-600 ng); uDNA1, non-specific DNA1 (*HacPGK2*); uDNA2, non-specific DNA2 (*HaCWI3*). The sunflower KARs are named according to previous publications and they are followed by the chromosome number where they are located.

**Suppl. Table 1. Oligonucleotides used in this work.**

**Suppl. Table 2. Promoter regions of the sunflower FAT and SAD genes described in literature.** Oligonucleotide pairs for PCR cloning, product length (bp), position relative to the TIS (bp) and presence of a AW box motif (YES or NO): FAT, Acyl-ACP thioesterase; SAD, Stearoyl-ACP desaturase; TIS, Translational initiation site.

**Suppl. Table 3. EMSA DNA fragments from sunflower FAT and SAD promoter regions and control genes.** Oligonucleotide pairs for PCR, PCR product length (bp) and location, and the position of the AW box motif relative to the TIS (bp). FAT, Acyl-ACP thioesterase; SAD, Stearoyl-ACP desaturase; cPGK, cytosolic phosphoglycerate kinase; CWI3, cell wall invertase 3; TIS, Translational Initiation Site; uDNA, non-specific DNA; N.A., non-applicable.

**Suppl. Table 4. EMSA DNA fragments from upstream regions of sunflower genes studied here that are involved in intraplastidial fatty acid biosynthesis.** The Heliagene ID-v2020 is indicated for each gene sequence together with the gene name stated in this work, based on the gene’s function and chromosome location: *Heliagene ID-v2018. The oligonucleotide pairs, EMSA PCR product length (bp) and position relative to the TIS (bp) are indicated for each fragment analyzed. The position of the AW box is indicated as initial/end bp from the TIS, except for the EMSA DNA fragments that contain more than one motif where only the initial base is indicated for each. PDH, Pyruvate dehydrogenase; DHLAT, Dihydrolipoamide acetyl transferase; LPD, Dihydrolipoamide dehydrogenase; CT, Carboxyltransferase; BC, Biotin carboxylase; BCCP, Biotin carboxyl carrier protein; KAR, β-ketoacyl-ACP reductase; ENR, Enoyl-ACP reductase; KAS, Ketoacyl-ACP synthase; ACP, Acyl carrier protein; FAT, Acyl-ACP thioesterase; TIS, Translational Initiation Site.

**Suppl. Table 5. EMSA DNA fragments from sunflower genes obtained by site-directed mutagenesis overlap extension PCR.** Sunflower gene names are indicated according to their function and chromosome location. The position of the AW box motif is indicated relative to the initial/end bp of the TIS. The oligonucleotide pairs used for sequential PCR, the PCR product lengths, and the characteristics and length of the final EMSA DNAs are indicated for each fragment studied. LPD, Dihydrolipoamide dehydrogenase; KAR, β-ketoacyl-ACP reductase; ACP, Acyl carrier protein; FAT, Acyl-ACP thioesterase; TIS, Translational Initiation Site.

## References

Almagro-Armenteros, J.J., Sønderby, C.K., Sønderby S.K., Nielsen H., and Winther O. (2017) DeepLoc: prediction of protein subcellular localization using deep learning. Bioinformatics 33, 3387–3395. doi: 10.1093/bioinformatics/btx431

An, D., Kim, H., Ju, S., Go, Y.S., Kim, H.U., and Suh, M.C. (2017) Expression of *Camelina* WRINKLED1 isoforms rescue the seed phenotype of the *Arabidopsis wri1* mutant and increase the triacylglycerol content in tobacco leaves. Front. Plant Sci. 8:34. doi: 10.3389/fpls.2017.00034

Aznar-Moreno, J.A., Sánchez, R., Gidda, S.K., Martínez-Force, E., Moreno-Pérez, A.J., Venegas-Calerón, M., Garcés, R., Mullen, R.T., and Salas, J.J. (2018) New insights into sunflower (*Helianthus annuus* L.) FatA and FatB thioesterases, their regulation, structure and distribution. Front. Plant Sci. 9, 1496. doi: 10.3389/fpls.2018.01496

Aznar-Moreno, J.A., Venegas-Calerón, M., Du, Z.Y., Garcés, R., Tanner, J.A., Chye, M.L., Martínez-Force, E., and Salas, J.J. (2016a) Characterization of a small acyl-CoA-binding protein (ACBP) from *Helianthus annuus* L. and its binding affinities. Plant Physiol. Biochem. 102, 141–150. doi: 10.1016/j.plaphy.2016.02.025

Aznar-Moreno, J.A., Venegas-Calerón, M., Du, Z.Y., Garcés, R., Tanner, J.A., Chye, M.L., Martínez-Force, E., and Salas, J.J. (2020) Characterization and function of a sunflower (*Helianthus annuus* L.) Class II acyl-CoA-binding protein. Plant Sci. 300, 110630. doi: 10.1016/j.plantsci.2020.110630

Aznar-Moreno, J.A., Venegas-Calerón, M., Martínez-Force, E., Garcés, R., and Salas, J.J. (2016b) Acyl carrier proteins from sunflower (*Helianthus annuus* L.) seeds and their influence on FatA and FatB acyl-ACP thioesterase activities. Planta 244, 479–490. doi: 10.1007/s00425-016-2521-7

Badouin, H., Gouzy, J., Grassa, C.J., Murat, F., Staton, S.E., Cottret, L., Lelandais-Brière, C., Owens, G.L., Carrère, S., Mayjonade, B., Legrand, L., Gill, N., Kane, N.C., Bowers, J.E., Hubner, S., Bellec, A., Bérard, A., Bergès, H., Blanchet, N., Boniface, M-C., Brunel, D., Catrice, O., Chaidir, N., Claudel, C., Donnadieu, C., Faraut, T., Fievet, G., Helmstetter, N., King, M., Knapp, S.J., Lai, Z., Le Paslier, M-C., Lippi, Y., Lorenzon, L., Mandel, J.R., Marage, G., Marchand, G., Marquand, E., Bret-Mestries, E., Morien, E., Nambeesan, S., Nguyen, T., Pegot-Espagnet, P., Pouilly, N., Raftis, F., Sallet, E., Schiex, T., Thomas, J., Vandecasteele, C., Varès, D., Vear, F., Vautrin, S., Crespi, M., Mangin, B., Burke, J.M., Salse, J., Muños, S., Vincourt, P., Rieseberg, L.H., and Langlade, N.B. (2017) The sunflower genome provides insights into oil metabolism, flowering and Asterid evolution. Nature 546:148–152. doi: 10.1038/nature22380

Bailey, T.L., and Gribskov, M. (1998) Combining evidence using p-values: application to sequence homology searches. Bioinformatics 14: 48–54. doi: 10.1093/bioinformatics/14.1.48

Baud, S., and Lepiniec, L. (2009) Regulation of de novo fatty acid synthesis in maturing oilseeds of Arabidopsis. Plant Physiol. Biochem. 47, 448–455. doi: 10.1016/j.plaphy.2008.12.006

Baud, S., and Lepiniec, L. (2010) Physiological and developmental regulation of seed oil production. Prog. Lipid Res. 49, 235–249. doi: 10.1016/j.plipres.2010.01.001

Baud, S., Mendoza, M.S., To, A., Harscoet, E., Lepiniec, L., and Dubreucq, B. (2007) WRINKLED1 specifies the regulatory action of LEAFY COTYLEDON2 towards fatty acid metabolism during seed maturation in Arabidopsis. Plant J. 50, 825–838. doi: 10.1111/j.1365-313X.2007.03092.x

Baud, S., Wuillème, S., To, A., Rochat, C., and Lepiniec, L. (2009) Role of WRINKLED1 in the transcriptional regulation of glycolytic and fatty acid biosynthetic genes in Arabidopsis. Plant J. 60: 933–947. doi: 10.1111/j.1365-313X.2009.04011.x

Berardini, T.Z., Reiser, L., Li, D., Mezheritsky, Y., Muller, R., Strait, E., and Huala, E. (2015) The Arabidopsis Information Resource: Making and mining the “gold standard” annotated reference plant genome. Genesis 53:474–485. doi: 10.1002/dvg.22877

Braybrook, S.A., and Harada, J.J. (2008) LECs go crazy in embryo development. Trends Plant Sci. 13, 624–630. doi: 10.1016/j.tplants.2008.09.008

Cernac, A., and Benning, C. (2004) WRINKLED1 encodes an AP2/EREB domain protein involved in the control of storage compound biosynthesis in Arabidopsis. Plant J. 40, 575–585. doi: 10.1111/j.1365-313X.2004.02235.x

Chang, W.C., Lee, T.Y., Huang, H.D., Huang, H.Y., and Pan, R.L. (2008) PlantPAN: Plant promoter analysis navigator, for identifying combinatorial cis-regulatory elements with distance constraint in plant gene groups. BMC Genomics 9: 561. doi: 10.1186/1471-2164-9-561

Chapman, K.D., and Ohlrogge, J.B. (2012) Compartmentation of triacylglycerol accumulation in plants. J. Biol. Chem. 287, 2288–2294. doi: 10.1074/jbc.R111.290072

Chen, B.B., Zhang, G.Y., Li, P.H., Yang, J.H., Guo, L., Benning, C., Wang, X.M., and Zhao, J. (2020) Multiple GmWRI1s are redundantly involved in seed filling and nodulation by regulating plastidic glycolysis, lipid biosynthesis and hormone signalling in soybean (*Glycine max*). Plant Biotechnol. J. 18, 155–171. doi: 10.1111/pbi.13183

Crooks G.E., Hon G., Chandonia J.M., and Brenner S.E. (2004) WebLogo: a sequence logo generator. Genome Res. 14, 1188–1190. doi: 10.1101/gr.849004

Durrett, T.P., Benning, C., and Ohlrogge, J. (2008) Plant triacylglycerols as feedstocks for the production of biofuels. Plant J. 54, 593–607. 10.1111/j.1365-313X.2008.03442.x

Fei, W., Yang, S., Hu, J., Yang, F., Qu, G., Peng, D., and Zhou, B. (2020) Research advances of WRINKLED1 (WRI1) in plants. Funct. Plant Biol. 47, 185–194. doi: 10.1071/FP19225

Focks, N., and Benning, C. (1998) *Wrinkled1*: A novel, low-seed-oil mutant of Arabidopsis with a deficiency in the seed-specific regulation of carbohydrate metabolism. Plant Physiol. 118, 91–101. doi: 10.1104/pp.118.1.91

Fukuda, N., Ikawa, Y., Aoyagi, T., and Kozaki, A. (2013) Expression of the genes coding for plastidic acetyl-CoA carboxylase subunits is regulated by a location-sensitive transcription factor binding site. Plant Mol. Biol. 82, 473–483. doi: 10.1007/s11103-013-0075-7

Gill, N., Buti, M., Kane, N., Bella, A., Helmsletter, N., Berges, H., and Rieseberg, L.H. (2014) Sequence-based analysis of structural organization and composition of the cultivated sunflower (*Helianthus annuus* L.) genome. Biology 3, 295–319. doi: 10.3390/biology3020295

Girke, T., Todd, J., Ruuska, S., White, J., Benning, C., and Ohlrogge, J. (2000) Microarray analysis of developing Arabidopsis seeds. Plant Physiol. 124, 1570–1581. doi: 10.1104/pp.124.4.1570

González-Mellado, D., von Wettstein-Knowles, P., Garcés, R., and Martínez-Force, E. (2010) The role of beta-ketoacyl-acyl carrier protein synthase III in the condensation steps of fatty acid biosynthesis in sunflower. Planta 231: 1277–1289. doi: 10.1007/s00425-010-1131-z

González-Mellado, D., Salas, J.J., Venegas-Calerón, M., Moreno-Pérez, A.J., Garcés, R., and Martínez-Force, E. (2019) Functional characterization and structural modelling of *Helianthus annuus* (sunflower) ketoacyl-CoA synthases and role in seed oil composition. Planta 249, 1823–1836. doi: 10.1007/s00425-019-03126-1

González-Thuillier, I., Venegas-Calerón, M., Moreno-Pérez, A.J., Salas, J.J., Garcés, R., von Wettstein-Knowles, P., and Martínez-Force, E. (2021) Sunflower (*Helianthus annuus*) fatty acid synthase complex: β-Ketoacyl-[acyl carrier protein] reductase genes. Plant Physiol. Biochem. 166, 689–699. doi: 10.1016/j.plaphy.2021.06.048

González-Thuillier, I., Venegas-Calerón, M., Garcés, R., von Wettstein-Knowles, P., and Martínez-Force, E. (2015) Sunflower (*Helianthus annuus*) fatty acid synthase complex: Enoyl-[acyl carrier protein]-reductase genes. Planta 241, 43–56. doi: 10.1007/s00425-014-2162-7

González-Thuillier, I., Venegas-Calerón, M., Sánchez, R., Garcés, R., von Wettstein-Knowles, P., and Martínez-Force, E. (2016) Sunflower (*Helianthus annuus*) fatty acid synthase complex: β-hydroxyacyl-[acyl carrier protein] dehydratase genes. Planta 243, 397–410. doi: 10.1007/s00425-015-2410-5

Heckman, K.L., and Pease, L.R. (2007) Gene splicing and mutagenesis by PCR-driven overlap extension. Nat. Protoc. 2, 924–932. doi: 10.1038/nprot.2007.132

Hongtrakul, V., Slabaugh, M.B., and Knapp, S.J. (1998) DFLP, SSCP, and SSR markers for delta 9-stearoyl-acyl carrier protein desaturases strongly expressed in developing seeds of sunflower: intron lengths are polymorphic among elite inbred lines. Mol. Breed. 4, 195–203. doi: 10.1023/A:1009646720400

Ji, X.J., Mao, X., Hao, Q.T., Liu, B.L., Xue, J.A., and Li, R.Z. (2018) Splice variants of the castor *WRI1* gene upregulate fatty acid and oil biosynthesis when expressed in tobacco leaves. Int. J. Mol. Sci. 19, 146. doi: 10.3390/ijms19010146

Kazaz, S., Barthole, G., Domergue, F., Ettaki, H., To, A., Vasselon, D., De Vos, D., Belcram, K., Lepiniec, L., and Baud, S. (2020) Differential activation of partially redundant Δ9 stearoyl-ACP desaturase genes is critical for omega-9 monounsaturated fatty acid biosynthesis during seed development in Arabidopsis. Plant Cell 32, 3613–3637. doi: 10.1105/tpc.20.00554

Kong, Q., Yang, Y.Z., Guo, L., Yuan, L., and Ma, W. (2020) Molecular basis of plant oil biosynthesis: Insights gained from studying the WRINKLED1 transcription factor. Front. Plant Sci. 11, 24. doi: 10.3389/fpls.2020.00024

Krizek, B.A. (2003) AINTEGUMENTA utilizes a mode of DNA recognition distinct from that used by proteins containing a single AP2 domain. Nucleic Acids Res. 31, 1859–1868. doi: 10.1093/nar/gkg292

Kuczynski, C., McCorkle, S., Keereetaweep, J., Shanklin, J., and Schwender J. (2020) An expanded role for WRINKLED1 metabolic control based on combined phylogenetic and biochemical analyses. bioRxiv doi: 10.1101/2020.01.28.923292

Li, Q., Shao, J., Tang, S., Shen, Q., Wang, T., Chen, W., and Hong, Y. (2015) Wrinkled1 accelerates flowering and regulates lipid homeostasis between oil accumulation and membrane lipid anabolism in *Brassica napus*. Front. Plant Sci. 6, 1015. doi: 10.3389/fpls.2015.01015. Erratum in: Front. Plant Sci. 6, 1270. doi: 10.3389/fpls.2015.01270

Li-Beisson, Y., Shorrosh, B., Beisson, F., Andersson, M.X., Arondel, V., Bates, P.D., Baud, S., Bird, D., Debono, A., Durrett, T.P., Franke, R.B., Graham, I.A., Katayama, K., Kelly, A.A., Larson, T., Markham, J.E., Miquel, M., Molina, I., Nishida, I., Rowland, O., Samuels, L., Schmid, K.M., Wada, H., Welti, R., Xu, C., Zallot, R., and Ohlrogge, J. (2013) “Acyl-lipid metabolism” in Arabidopsis Book Vol 11, e0161. doi: 10.1199/tab.0161

Liu, H., Zhai, Z., Kuczynski, K., Keereetaweep, J., Schwender, J., and Shanklin, J. (2019) WRINKLED1 regulates Biotin Attachment Domain-Containing proteins that inhibit fatty acid synthesis. Plant Physiol. 181, 55–62. doi: 10.1104/pp.19.00587

Liu, J., Hua, W., Zhan, G.M., Wei, F., Wang, X.F., Liu, G.H., and Wang, H.Z. (2010) Increasing seed mass and oil content in transgenic Arabidopsis by the overexpression of wri1-like gene from *Brassica napus*. Plant Physiol. Biochem. 48, 9–15. doi: 10.1016/j.plaphy.2009.09.007

Ma, W., Kong, Q., Arondel, V., Kilaru, A., Bates, P.D., Thrower, N.A., Benning, C., and Ohlrogge, J.B. (2013) *WRINKLED1*, a ubiquitous regulator in oil accumulating tissues from *Arabidopsis* embryos to oil Palm mesocarp. PLoS One 8, e68887. Doi: 10.1371/journal.pone.0068887

Maeo, K., Tokuda, T., Ayame, A., Mitsui, N., Kawai, T., Tsukagoshi, H., Ishiguro, S., and Nakamura, K. (2009) An AP2-type transcription factor, WRINKLED1, of Arabidopsis thaliana binds to the AW-box sequence conserved among proximal upstream regions of genes involved in fatty acid synthesis. Plant J. 60, 476–487. doi: 10.1111/j.1365-313X.2009.03967.x

Mano, F., Aoyanagi, T., and Kozaki, A. (2019) Atypical splicing accompanied by skipping conserved micro-exons produces unique WRINKLED1, an AP2 domain transcription factor in rice plants. Plants 8, 207. doi: 10.3390/plants8070207

Martins-Noguerol, R., Moreno-Pérez, A.J., Acket, S., Makni, S., Garcés, R., Troncoso-Ponce, A., Salas, J.J., Thomasset, B., and Martínez-Force, E. (2019) Lipidomic analysis of plastidial octanoyltransferase mutants of *Arabidopsis thaliana*. Metabolites 9, 209. doi: 10.3390/metabo9100209

Martins-Noguerol, R., Moreno-Pérez, A.J., Acket, S., Troncoso-Ponce, M.A., Garcés, R., Thomasset, B., Salas, J.J., and Martínez-Force, E. (2020) Impact of sunflower (*Helianthus annuus* L.) plastidial lipoyl synthases genes expression in glycerolipids composition of transgenic Arabidopsis plants. Sci. Rep. 10, 3749. doi: 10.1038/s41598-020-60686-z

Mentzen, W.I., Peng, J.L., Ransom, N., Nikolau, B.J., and Wurtele, E.S. (2008) Articulation of three core metabolic processes in Arabidopsis: Fatty acid biosynthesis, leucine catabolism and starch metabolism. BMC Plant Biol. 8, 76. doi: 10.1186/1471-2229-8-76

Miray, R., Kazaz, S., To, A., and Baud, S. (2021) Molecular control of oil metabolism in the endosperm of seeds. Int. J. Mol. Sci. 22, 1621. doi: 10.3390/ijms22041621

Moreno-Pérez, J.A., Santos-Pereira, J.M., Martins-Noguerol, R., DeAndrés-Gil, C., Troncoso-Ponce, M.A., Venegas-Calerón, M., Sánchez, R., Garcés, R., Salas, J.J., Tena, J.J., and Martínez-Force, E. (2021) Genome-wide mapping of histone H3 lysine 4 trimethylation (H3K4me3) and its involvement in fatty acid biosynthesis in sunflower developing seeds. Plants 10, 706. doi: 10.3390/plants10040706

North, H., Baud, S., Debeaujon, I., Dubos, C., Dubreucq, B., Grappin, P., Jullien, M., Lepiniec, L., Marion-Poll, A., Miquel, M., Rajjou, L., Routaboul, J.M., and Caboche, M. (2010) Arabidopsis seed secrets unravelled after a decade of genetic and omics-driven research. Plant J. 61, 971–981. doi: 10.1111/j.1365-313X.2009.04095.x

Ohlrogge, J., and Chapman, K. (2011) The seeds of green energy: expanding the contribution of plant oils as biofuels. Biochemist 33, 34–38. doi: 10.1042/BIO03302034

Penouilh-Suzette, C., Pomiès, L., Duruflé, H., Blanchet, N., Bonnafous, F., Dinis, R., Brouard, C., Gody, L., Grassa, C., Heudelot, X., Laporte, M., Larroque, M., Marage, G., Mayjonade, B., Mangin, B., de Givry, S., and Langlade N.B. (2020) RNA expression dataset of 384 sunflower hybrids in field condition. OCL 27, 36. doi: 10.1051/ocl/2020027

Pouvreau, B., Baud, S., Vernoud, V., Morin, V., Py, C., Gendrot, G., Pichon, J-P., Rouster, J., Paul, W., and Rogowsky, P.M. (2011) Duplicate maize wrinkled1 transcription factors activate target genes involved in seed oil biosynthesis. Plant Physiol. 156: 674–686. doi: 10.1104/pp.111.173641

Ruuska, S.A., Girke, T, Benning, C., and Ohlrogge, J.B. (2002) Contrapuntal networks of gene expression during Arabidopsis seed filling. Plant Cell 14, 1191–1206. doi: 10.1105/tpc.000877

Salas, J.J., Martínez-Force, E., Harwood, J.L., Venegas-Calerón, M., Aznar-Moreno, J.A., Moreno-Pérez, A.J., Ruíz-López, N., Serrano-Vega, M.J., Graham, I.A., Mullen, R.T., and Garcés, R. (2014) Biochemistry of high stearic sunflower, a new source of saturated fats. Prog. Lipid Res. 55, 30–42. doi: 10.1016/j.plipres.2014.05.001

Schmid, M., Davison, T.S., Henz, S.R., Pape, U.J., Demar, M., Vingron, M., Schölkopf, B., Weigel, D., and Lohmann, J.U. (2005) A gene expression map of *Arabidopsis thaliana* development. Nat. Genet. 37, 501–506. doi: 10.1038/ng1543

Serrano-Vega, M.J., Venegas-Calerón, M., Garcés, R., and Martínez-Force, E. (2003) Cloning and expression of fatty acids biosynthesis key enzymes from sunflower (*Helianthus annuus* L.) in *Escherichia coli*. J. Chromatogr. B Analyt. Technol. Biomed. Life Sci. 786, 221–228. doi: 10.1016/s1570-0232(02)00767-5

Serrano-Vega, M.J., Garcés, R., and Martínez-Force, E. (2005) Cloning, characterization and structural model of a FatA-type thioesterase from sunflower seeds (*Helianthus annuus* L.). Planta 221, 868–880. doi: 10.1007/s00425-005-1502-z

Shen, B., Allen, W.B., Zheng, P.Z., Li, C.J., Glassman, K., Ranch, J., Nubel, D., and Tarczynski, M.C. (2010) Expression of ZmLEC1 and ZmWRI1 increases seed oil production in maize. Plant Physiol. 153, 980–987. doi: 10.1104/pp.110.157537

Sperschneider, J., Catanzariti, A-M., DeBoer, K., Petre, B., Gardiner, D.M., Singh, K.B., Dodds, P.N., and Taylor, J.M. (2017) LOCALIZER: subcellular localization prediction of both plant and effector proteins in the plant cell. Sci. Rep. 7, 44598. doi: 10.1038/srep44598

Suzuki, M., and McCarty, D.R. (2008) Functional symmetry of the B3 network controlling seed development. Curr. Opin. Plant Biol. 11, 548–553. doi: 10.1016/j.pbi.2008.06.015

Tajima, D., Kaneko, A., Sakamoto, M., Ito, Y., Hue, N., Miyazaki, M., Ishibashi, Y., Yuasa, T., and Iwaya-Inoue, M. (2013) *Wrinkled* 1 (WRI1) homologs, AP2-type transcription factors involving master regulation of seed storage oil synthesis in castor bean (*Ricinus communis* L.). Am. J. Plant Sci. 4, 333–339. doi: 10.4236/ajps.2013.42044

To, A., Joubès, J., Barthole, G., Lécureuil, A., Scagnelli, A., Jasinski, S., Lepiniec, L., and Baud, S. (2012) WRINKLED transcription factors orchestrate tissue-specific regulation of fatty acid biosynthesis in Arabidopsis. Plant Cell 24: 5007–5023. doi: 10.1105/tpc.112.106120

Venegas-Calerón, M., Troncoso-Ponce, M.A., and Martínez-Force, E. (2015) “Sunflower Oil and Lipids Biosynthesis”, in Sunflower Oilseed. Chemistry, Production, Processing and Utilization, AOCS Monograph Series on Oilseeds Vol. 7, eds. E. Martínez-Force, N.T. Dunford and J. J. Salas (Urbana, IL: AOCS Press), 259–295. doi: 10.1016/B978-1-893997-94-3.50016-7

Yang, Y., Munz, J., Cass, C., Zienkiewicz, A., Kong, Q., Ma, W., Sanjaya, Sedbrook, J., and Benning, C. (2015) Ectopic expression of WRINKLED1 affects fatty acid homeostasis in *Brachypodium distachyon* vegetative tissues. Plant Physiol. 169, 1836–1847. doi: 10.1104/pp.15.01236

Zambelli, A., León, A., and Garcés, R. (2015) “Mutagenesis in Sunflower”, in Sunflower Oilseed. Chemistry, Production, Processing and Utilization, AOCS Monograph Series on Oilseeds Vol. 7, eds. E. Martínez-Force, N.T. Dunford and J. J. Salas (Urbana, IL: AOCS Press), 27–52. doi: 10.1016/B978-1-893997-94-3.50008-8

Zeinalzadehtabrizi, H., Hosseinpour, A., Aydin, M., and Haliloglu, K. (2015) A modified genomic DNA extraction method from leaves of sunflower for PCR based analyzes. J. Biodivers. Environ. Sci. 7: 222–225

