## Supplementary figures and images for "The sunflower WRINKLED1 transcription factor regulates fatty acid biosynthesis genes through an AW box binding sequence with a particular base bias"

### Supplemental Fig1

Supplementary Figure 1

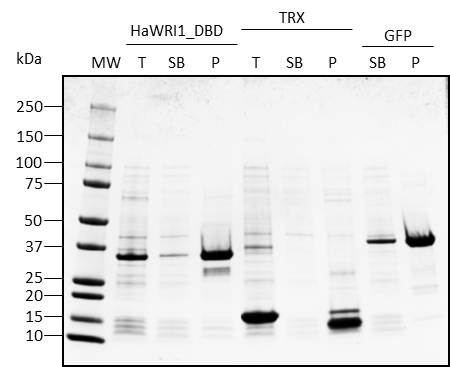

### Supplemental Fig2

Supplementary Figure 2

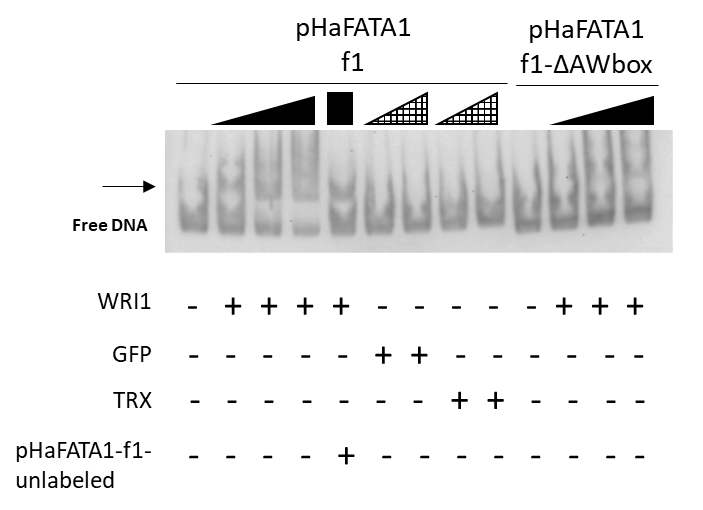

### Supplemental Fig3

Supplementary Figure 3

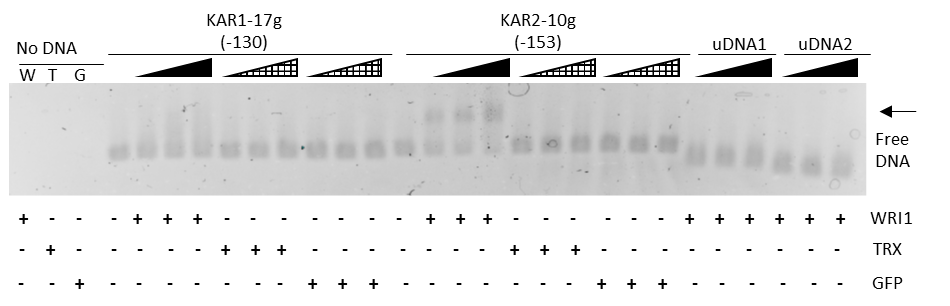
